# Concerted but segregated actions of oxytocin and vasopressin within the ventral and dorsal lateral septum determine female aggression

**DOI:** 10.1101/2020.07.28.224873

**Authors:** Vinícius Elias de Moura Oliveira, Michael Lukas, Hannah Nora Wolf, Elisa Durante, Alexandra Lorenz, Anna-Lena Mayer, Anna Bludau, Oliver J. Bosch, Valery Grinevich, Veronica Egger, Trynke R. de Jong, Inga D. Neumann

## Abstract

In contrast to males, aggression in females has been rarely studied. Here, we established a rat model of enhanced female aggression using a combination of social isolation and aggression-training to specifically investigate the involvement of the oxytocin (OXT) and vasopressin (AVP) systems within the lateral septum (LS). Using neuropharmacological, optogenetic, chemogenetic as well as microdialyses approaches, we revealed that enhanced OXT release within the ventral LS (vLS), combined with reduced AVP release within the dorsal LS (dLS), are required for female aggression. Accordingly, increased excitability of OXT-responsive neurons in the vLS and decreased excitability of AVP-responsive neurons in the dLS were essential to evoke female aggression. Finally, *in vitro* activation of OXT receptors in the vLS increased tonic GABAergic inhibition of dLS neurons. Overall, our data demonstrate that septal release of OXT and AVP affects female aggression by differential regulation of the excitatory-inhibitory balance within subnetworks of the LS.

## Introduction

Aggressive behavior is expressed by most, if not all, mammalian species, including humans. Typically, aggression is displayed whenever conspecifics have to compete for resources, such as food, territory, or mating partners^1^. When expressed out-of-context or in an exacerbated manner, aggressive behavior becomes disruptive and harms both aggressor and victim. In humans, excessive or pathological aggression as seen, for example, in individuals suffering from conduct or antisocial personality disorder constitutes a severe burden to society^1,2^. To better understand the neurobiology of aggression and to develop potential treatment options, laboratory animal models of aggression have been successfully used for decades^1,3,4^. However, these models were mostly developed in male rodents, whereas females have been rather understudied, except during the physiologically unique period of lactation^5^. Given the fact that i) girls and women also demonstrate disruptive aggression and may develop conduct as well as antisocial personality disorder similarly to males^6–8^, and ii) the neurobiology of aggression appears to be sex-dimorphic^5,8–11^, the use of non-lactating female animal models is required to increase our understanding of the neurobiology of female aggression and to identify potential targets for the treatment of pathological aggression in both sexes.

In order to study the neurobiological mechanisms underlying female aggression, we first established an animal model to robustly enhance the mild levels of aggression displayed by virgin female Wistar rats^12^. We predicted that a combination of social isolation and aggression-training by repeated exposure to the female intruder test (FIT)^12^, i.e. to an unknown same-sex intruder, enhances female aggressiveness, since both social isolation^11,13^ and repeated engagement in conflict with conspecifics (“winner effect”)^14–16^, exacerbate aggression in solitary and aggressive rodent species, independent of sex.

Next, we investigated the role of oxytocin (OXT) and arginine vasopressin (AVP) in female aggression. Both neuropeptides have been associated with various social behaviors including aggression in males and lactating females^17–24^, and are known to be affected by social isolation in a sex-dependent manner^11,13^. In this context, we hypothesized that the effects of OXT and AVP on female aggression are predominantly mediated in the lateral septum (LS) - a target region for neuropeptide actions^24^, known to suppress aggression in both sexes^10,25^. Electrical stimulation of the LS^26,27^, and more specifically optogenetic stimulation of septal GABAergic projections to the ventromedial nucleus of the hypothalamus (VMH)^28^, reduces aggression in male rodents. In contrast, pharmacological inhibition or lesioning of the LS triggers exaggerated aggression (‘septal rage’) in male and female hamsters^10,29,30^. In support, male and female patients with septal tumors show increased irritability and aggressive outbursts^25^.

Although several pieces of evidence point towards the gating role of the LS in aggression, the underlying mechanisms, especially the involvement of septal OXT and AVP in female aggression, are still unknown. The release of both neuropeptides in the LS has been tied to various social behaviors, including intermale aggression, although the results are somewhat conflicting^24^. For example, in male Wistar rats attenuated^21^ as well as increased^22^ release of AVP have been described during the display of abnormal and high^31^ levels of aggression, respectively. Regarding the role of the septal OXT system, increased OXT receptor (OXTR) binding has been reported in both dominant male mice^32^ and lactating female rats^33^. Importantly, rats have a markedly specific expression of OXTR and AVP V1a receptors (V1aRs) in two distinct subregions of the LS, i.e. OXTRs are exclusively found in the ventral LS (vLS), whereas V1aRs are predominantly found in the dorsal LS (dLS)^34^.

Here, using the novel and reliable rat model of female aggression, we first compared central release patterns of endogenous OXT and AVP between low-aggressive (group-housed; GH) and high-aggressive (isolated and trained; IST) rats by measuring neuropeptide content in microdialysates sampled within the LS or in the cerebrospinal fluid (CSF), during or after the FIT, respectively. Additionally, we assessed septal OXTR and V1aR binding in GH and IST females. Next, we employed pharmacological, chemogenetic, and optogenetic approaches to selectively manipulate central OXT or AVP signaling, specifically in the vLS or dLS, and studied their behavioral consequences in the FIT. Finally, to specifically dissect the neuronal mechanisms within the LS controlling aggressive behavior, we activated septal OXTRs and V1aRs while recording spontaneous GABAergic inputs to the vLS and dLS GABAergic neurons *in vitro* and monitored as well as locally manipulated neuronal activity, in rats with opposite levels of aggression *in vivo*.

## Results

### Social isolation and training reliably enhance female aggression

Training consisted of daily 10-min exposure to a FIT^12^ on three consecutive days. GH and isolated non-trained rats (IS) were used as control groups to assess the effects of social isolation and aggression-training, respectively (Figure 1a). Both IST and IS females displayed heightened aggression, i.e. increased time spent on keep down, threat, and offensive grooming (Figure 1b), as well as spend more time and engaged more frequently in attacking (Figure 1c,d) compared to GH controls (Supplementary Movie1). We found a major effect of the estrus cycle on aggression, mainly reflected by lower levels of aggression in IS females in the proestrus or estrus phase than in the metestrus or diestrus phase (Figure 1e). Concerning other behaviors, GH females spent more time displaying non-social behaviors. Whereas only IST females showed increased self-grooming compared to GH females (Figure 1f).

**Figure 1.**
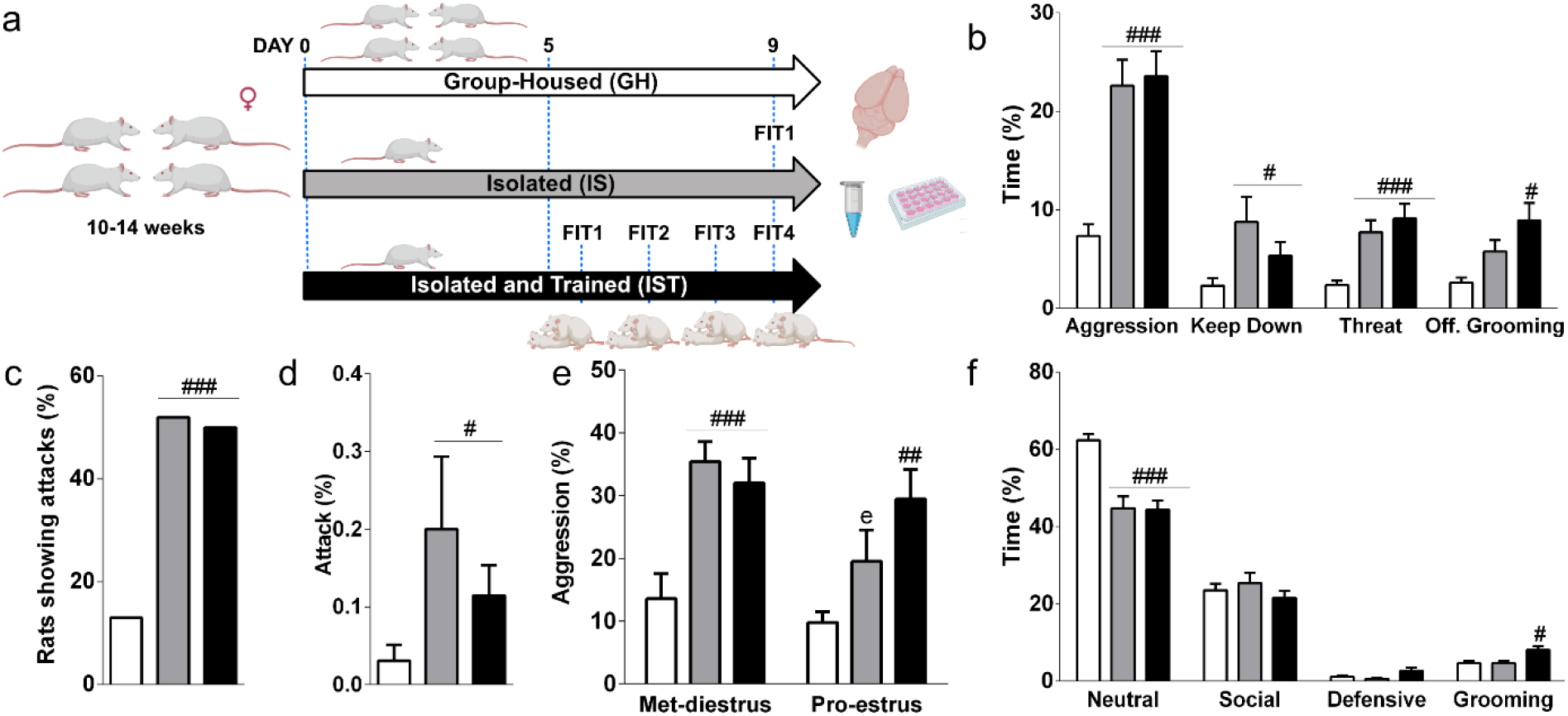
Social isolation and training reliably enhance female aggression, independently of the estrus cycle. **a** Scheme illustrating the animal groups and behavioral timeline. Female Wistar rats were kept either group-housed (GH) or socially-isolated (IS and IST). IST females were exposed to three consecutive female intruder tests (FIT1 to FIT3, training), i.e. to aggressive encounters with an unknown same-sex intruder. Partially drawn using https://biorender.com. **b** Both IS and IST rats showed increased total aggression (one-way ANOVA followed by Bonferroni F_(2,63)_=16.26, p< 0.0001), keep down (Kruskal-Wallis test followed by Dunn’s H_3_=6.69, p=0.035), threat (H_3_=18.98, p<0.0001) and offensive grooming (H_3_=10.8, p=0.005). **c** IS and IST females engaged more in attacks (Chi-squared test, X^2^=40.81, p<0.0001), and **d** spend more time attacking (H_3_=8.8, p=0.012) than GH females. **e** IS females in the proestrus and estrus (Pro-estrus) phase of the estrus cycle displayed less aggressive behavior than metestrus-diestrus (Met-diestrus) females (two-way ANOVA followed by Bonferroni; factor housing: F_(2, 57)_=14.2, p<0.0001; estrus cycle: F_(1, 57)_=5.47, p=0.023; housing x estrous cycle: F_(2, 57)_ =1.78, p=0.18). **f** IST and IS females compensated their increased aggression with decreased neutral behaviors (F_(2, 63)_=18.76, p<0.0001), only IST females spent more time self-grooming (F_(2, 63)_=5.79, p=0.005). All data are shown as mean+SEM. ^#^p<0.05, ^##^p<0.001, ^###^p<0.001 vs GH; ^e^p<0.05 vs met-diestrus. n=21-23.

To further validate our model of female aggression, we treated aggressive IST females with the selective serotonin reuptake inhibitor (SSRI) escitalopram (10mg/kg, s.c.), since SSRIs typically decrease aggression in rodents^35^. Indeed, escitalopram significantly reduced the time spent on aggression, i.e. on keeping down, threatening and attacking, and increased the latency to attack. Consequently, escitalopram-treated females displayed more defensive and neutral behaviors (Supplementary Figure 1).

### The display of high aggression in IST rats is accompanied by elevated OXT and decreased AVP release in the brain

IST, IS and GH rats were either exposed to the FIT or not (control) to assess the effects of housing conditions as well as an aggressive encounter on the endogenous OXT and AVP systems (Figure 1a). Elevated post-FIT OXT levels were found in the CSF of IST females compared with GH females (Figure 2a). Additionally, IST and IS rats exhibited higher levels of OXT in the CSF compared with their respective non-FIT control groups. Consistent with these findings, aggression levels were positively correlated with OXT content in the CSF, when all groups were combined (Figure 2b). Notably, plasma OXT concentrations did not follow the same pattern (Supplementary Figure 2a-b). In contrast to OXT, AVP concentrations in the CSF after FIT exposure were lower only in IST females without any effect of housing or training conditions (Figure 2f). Consequently, AVP content in the CSF did not correlate with aggression (Figure 2g). Regarding AVP secretion into the blood, FIT exposure tended to heighten plasma AVP regardless of housing condition, although the magnitude of this effect seemed to be reduced in aggressive IST females (Supplementary Figure 2d). Plasma AVP also did not correlate with aggression (Supplementary Figure 2e).

**Figure 2.**
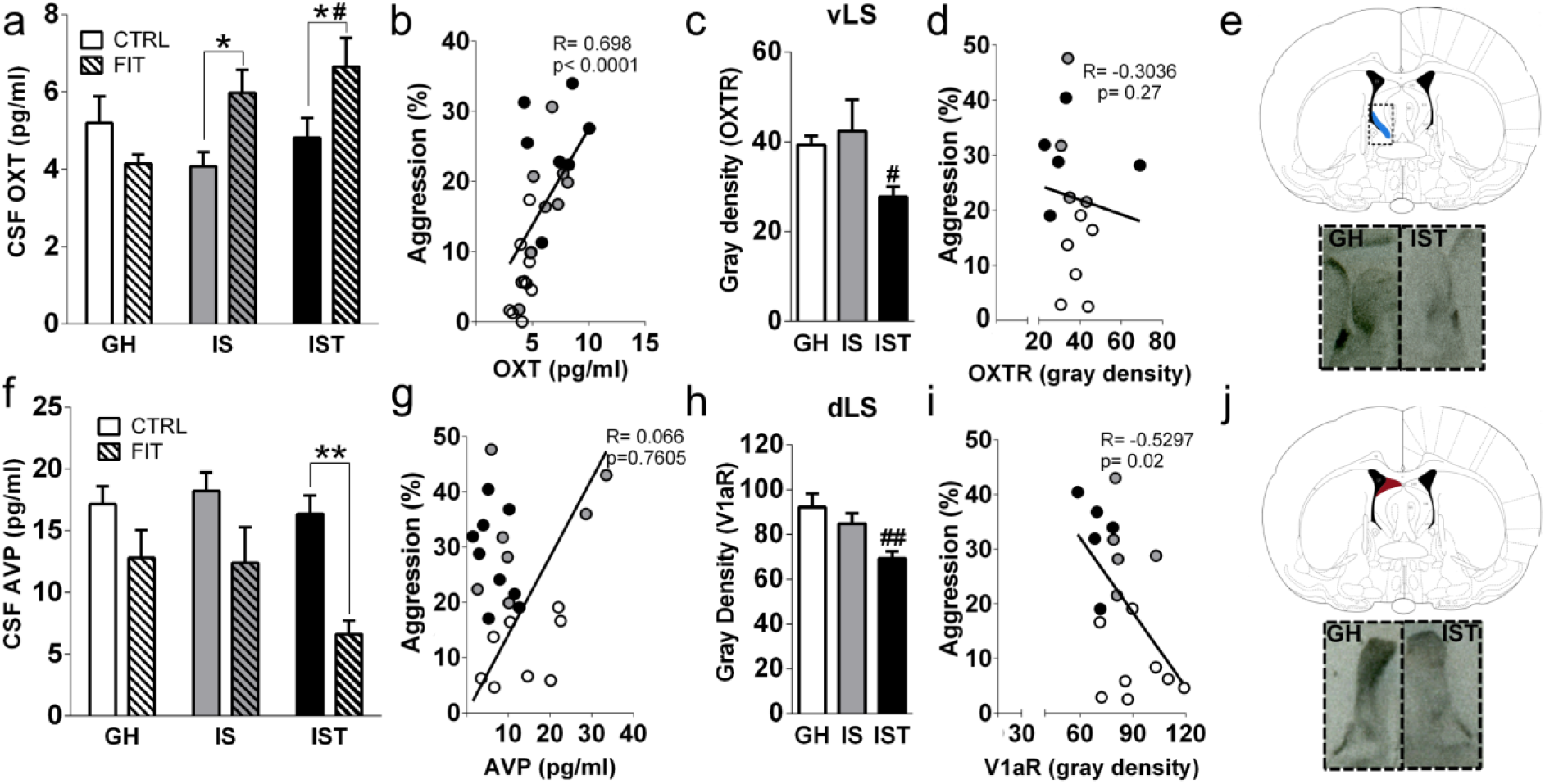
The high levels of aggression displayed by isolated and trained (IST) rats are accompanied by high OXT and low AVP concentrations in cerebrospinal fluid (CSF) and low OXT and V1a receptor binding in the lateral septum (LS). **a** IS and IST females showed higher concentrations of OXT in CSF immediately after exposure to the female intruder test (FIT) compared to respective control rats (CTRL) (two-way ANOVA followed by Bonferroni: factor FIT: F_(1,44)_=3.91, p=0.054; housing: F_(2,44)_=1.96, p=0.152; FIT x housing: F_(2,44)_=4.68, p=0.014). **b** Aggression correlated with CSF OXT levels of FIT rats (Pearson’s correlation r=0.70, p<0.0001). **c** IST females presented decreased OXT receptor (OXTR) binding in the ventral portion of the LS (vLS) (Kruskal-Wallis test followed by Dunn’s: H_(3)_=7.12, p=0.028), **d** which did not correlate with aggression levels (Spearman’s correlation r=−0.304, p=0.27). **e** Scheme illustrating localization of OXTR in the vLS, and magnification of representative example autoradiograph (left: GH; right: IST). **f** FIT exposure decreased CSF AVP levels only in IST rats (two-way ANOVA, factor FIT: F_(1,50)_=15.98, p=0.0002; housing: F_(2,50)_=2.13, p=0.129; FIT x housing: F_(2,50)_=0.90, p=0.41). **g** Aggressive behavior did not correlate with CSF AVP concentrations (r=0.066, p=0.76). **h** IST females presented decreased V1aR binding in the dorsal part of the LS (dLS) (H_(3)_=8.72, p=0.006). **i** Aggression negatively correlated with V1aR binding in the dLS (r=−0.53, p=0.02). **j** Scheme illustrating localization of V1aR in the dLS and magnification of representative autoradiograph (left: GH, right: IST). All data are presented as mean + SEM. ^#^p<0.05, ^##^p<0.01 vs GH; ^*^p<0.05, ^**^p<0.01 vs control. Binding: n=6-9; OXT: n=8-9; AVP: n=8-11;

### Highly aggressive IST rats had low OXTR and V1aR binding in the vLS and dLS, respectively

We analyzed OXTR and V1aR binding in selected brain regions involved in the social/aggressive behavior network (Supplementary Figure 2). Among the 9 regions analyzed, only within the LS OXTRs were affected by housing and training conditions, as IST females exhibited lower OXTR binding in the vLS compared to GH controls (Figure 2c and 2e). However, local OXTR binding did not correlate with aggression (Figure 2d). We also found lower V1aR binding within the dLS in IST females when compared with GH controls (Figure 2h and 2j, Supplementary Figure 2f), and a negative correlation of V1aR binding with aggression, i.e. female rats with reduced V1aR binding were found to be more aggressive (Figure 2i).

Since low glucocorticoid levels have been implicated in the development of intermale aggression in humans and animals ^1,8^, we assessed plasma corticosterone concentrations after exposure to the FIT. Although there was no effect of housing or training conditions, plasma corticosterone indeed negatively correlated with aggression in females (Supplementary figure 2g).

### OXT promotes, whereas AVP reduces female aggression

Because of the identified correlation between OXT content in the CSF and female aggression as well as the increased levels of OXT in highly aggressive females, we employed pharmacological and chemogenetic methods to either increase OXT availability in the brain of low-aggressive, i.e. GH females, or to block central OXT neurotransmission in the brain of high-aggressive, i.e. IST females (Figure 3a).

**Figure 3.**
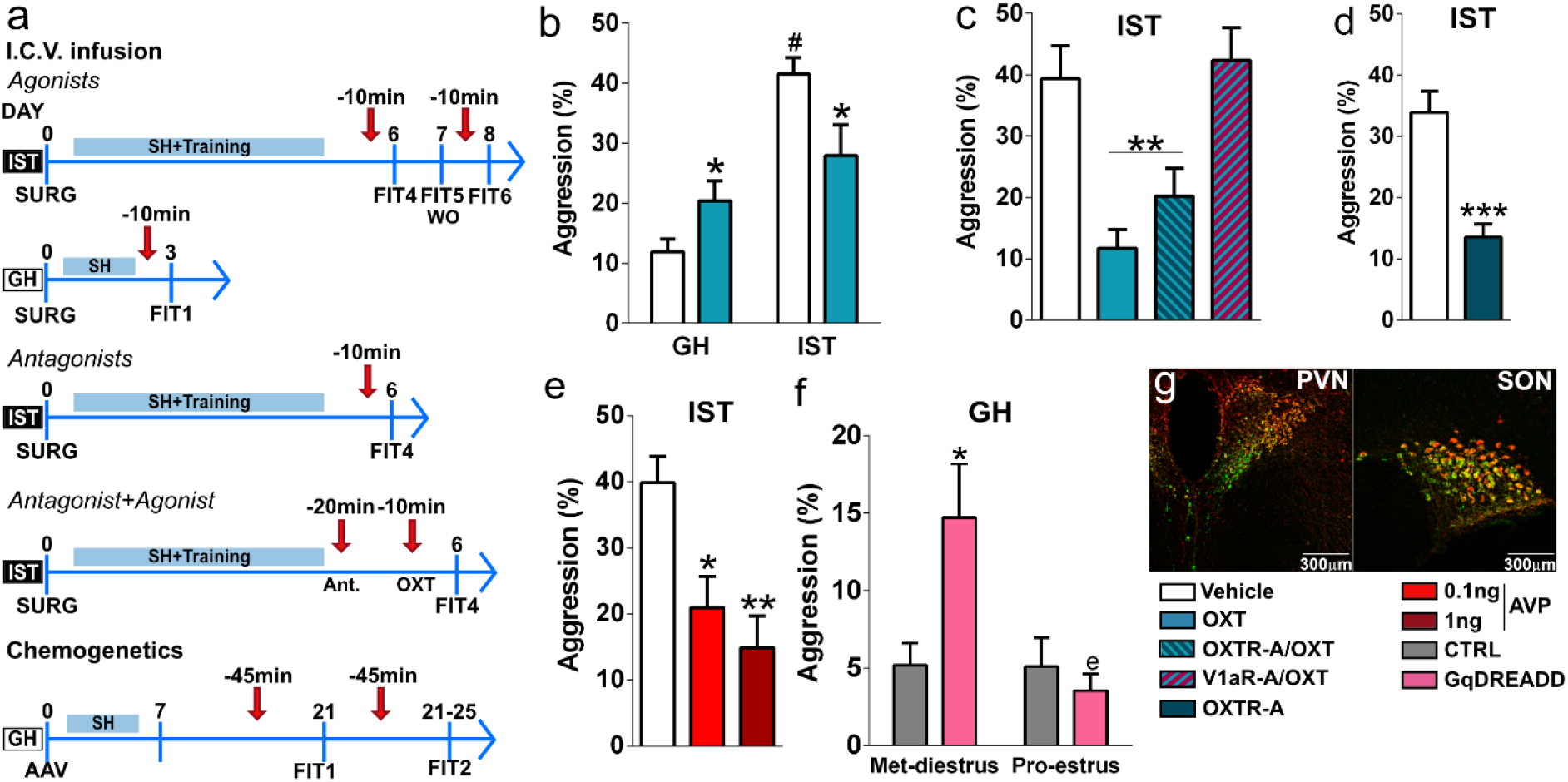
OXT promotes, whereas AVP reduces female aggression. **a** Experimental design for pharmacological and chemogenetic experiments targeting the OXT and AVP systems in isolated and trained (IST) and group-housed (GH) rats. (AAV= adeno-associated DREADD virus infusion into the hypothalamic paraventricular (PVN) and supraoptic (SON) nuclei; AVP= vasopressin; arrow= drug infusions; FIT= female intruder test; OXT= oxytocin; OXTR-A= OXT receptor antagonist; SH= single housing; SURG= surgery; V1aR-A= V1a receptor antagonist; WO= Wash-out). **b** I.c.v. infusion of OXT (50ng/5ul) increased aggression in GH (two-tailed Student’s t-test t_(19)_=2.46, p=0.024), but decreased aggression in IST females (t_(8)_=2.33, p=0.048). **c** I.c.v. infusion of V1aR-A (750ng/5μl), but not OXTR-A (750ng/5μl) blocked the anti-aggressive effects of OXT in IST females (one-way ANOVA followed by Bonferroni F_(3,28)_=10.1, p=0.001). Both, **d** i.c.v. infusion of OXTR-A (t_(28)_=4.96, p<0.0001) and **e** i.c.v. AVP (0.1 or 1ng/5μl) reduced total aggressive behavior (F_(3,54)_=7.48, p=0.0003) in IST rats. **f** Chemogenetic activation of OXT neurons in the PVN and SON increased aggression only in met-diestrus GH rats (two-way ANOVA, factor treatment: F_(1,19)_=3.342, p=0.083; estrus cycle: F_(1,19)_=6.68, p=0.018; treatment x estrus cycle: F_(1,19)_= 6.45, p=0.02). **g** Confirmation of virus infection in the PVN (right) and SON (left). OXT-neurophysin I staining: green; mCherry (virus): red. Scale bars 300μm. Data are shown as mean + SEM. ^#^p<0.05 vs GH; ^*^p<0.05; ^**^p<0.01; ^***^p<0.0001 vs either vehicle or control; ^e^p<0.05 vs met-diestrus. OXT: n=9-12; AVP: n=9-18; OXTR-A: n=8; Combination OXT/OXTR-A/V1aR-A: n= 7-9; Chemogenetics: n= 7-15.

In detail, intracerebroventricular (i.c.v.) infusion of *synthetic* OXT enhanced aggression in GH females. Surprisingly, the same treatment decreased aggression in IST females (Figure 3b), and this anti-aggressive effect of OXT was reflected in all behaviors analyzed (Supplementary Figure 3). Although these results might seem puzzling at first glance, the chemical similarity between OXT and AVP, which co-evolved from a single nonapeptide^36^, is known to result in cross-reactivity between both peptides and each other’s receptors *in vitro*^37^ and *in vivo*, including in the context of aggression^38^. To exclude cross-reactivity of OXT on V1aRs, we blocked either OXTR or V1aR with selective antagonists (OXTR-A and V1aR-A, i.c.v.)^37^ 10 min prior to i.c.v. OXT infusion. Whereas the OXTR-A did not abolish the anti-aggressive effects of OXT, pre-infusion with V1aR-A did (Figure 3c and Supplementary Figure 3), clearly indicating that the anti-aggressive effect of *synthetic* OXT in IST females is mediated via V1aRs.

In order to reveal the involvement of *endogenous* OXT in female aggression, we first blocked OXTRs in IST females and found that i.c.v. infusion of the OXTR-A before FIT exposure reduced total aggression and threat behavior (Figure 3d, Supplementary Figure 3). Aiming to chemogenetically stimulate intracerebral OXT release we infected the hypothalamic paraventricular (PVN) and supraoptic (SON) nuclei of GH female rats with an rAAV to selectively express GqDREADD under the control of an OXT promoter fragment (Figure 3g). The virus showed a high degree of cell-type specificity as 64.4% of cells in the PVN and 75.1% of cells in the SON were positive for both OXT and mCherry signals in accordance with previous data^39^. However, a few mCherry-positive cells (12±1.6 cells/per section, 118 cells in total) outside the PVN and SON devoid OXT immunosignals. Intraperitoneal application of the DREADD ligand clozapine-N-oxide dihydrochloride increased aggression in diestrous and metestrous GH females, thereby confirming our pharmacological results. Intriguingly, estrus, and proestrus GH females showed no increase in aggression. (Figure 3f and Supplementary figure 4).

Finally, since i.c.v. *synthetic* OXT decreased aggression via activation of V1aRs, and because low levels of AVP were found in the CSF of IST rats, we hypothesized that *synthetic* AVP i.c.v would also reduce aggression in IST females. Indeed, elevation of AVP availability prior to the FIT resulted in decreased total aggression, keeping down, threatening behaviors as well as time spent on attacks (Figure 3e and Supplementary Figure 3).

Taken together, these results demonstrate that the pro-aggressive effect of *endogenous* OXT is mediated via OXTRs, whereas the anti-aggressive effects of *synthetic* OXT and, particularly, *synthetic* AVP are mediated via V1aRs.

### The pro-aggressive effect of OXT is mediated in the vLS

Based on the fact that OXTR binding was exclusively reduced in the vLS of IST females (Figure 2c), and that the LS is known to regulate aggression^26–28^, we hypothesized that the pro-aggressive effect of *endogenous* OXT is mediated in the vLS. Thus, we used local *in vivo* microdialysis, neuropharmacological and optogenetic approaches to specifically study the role of the OXT system in the vLS in female aggression (Figure 4a)

**Figure 4.**
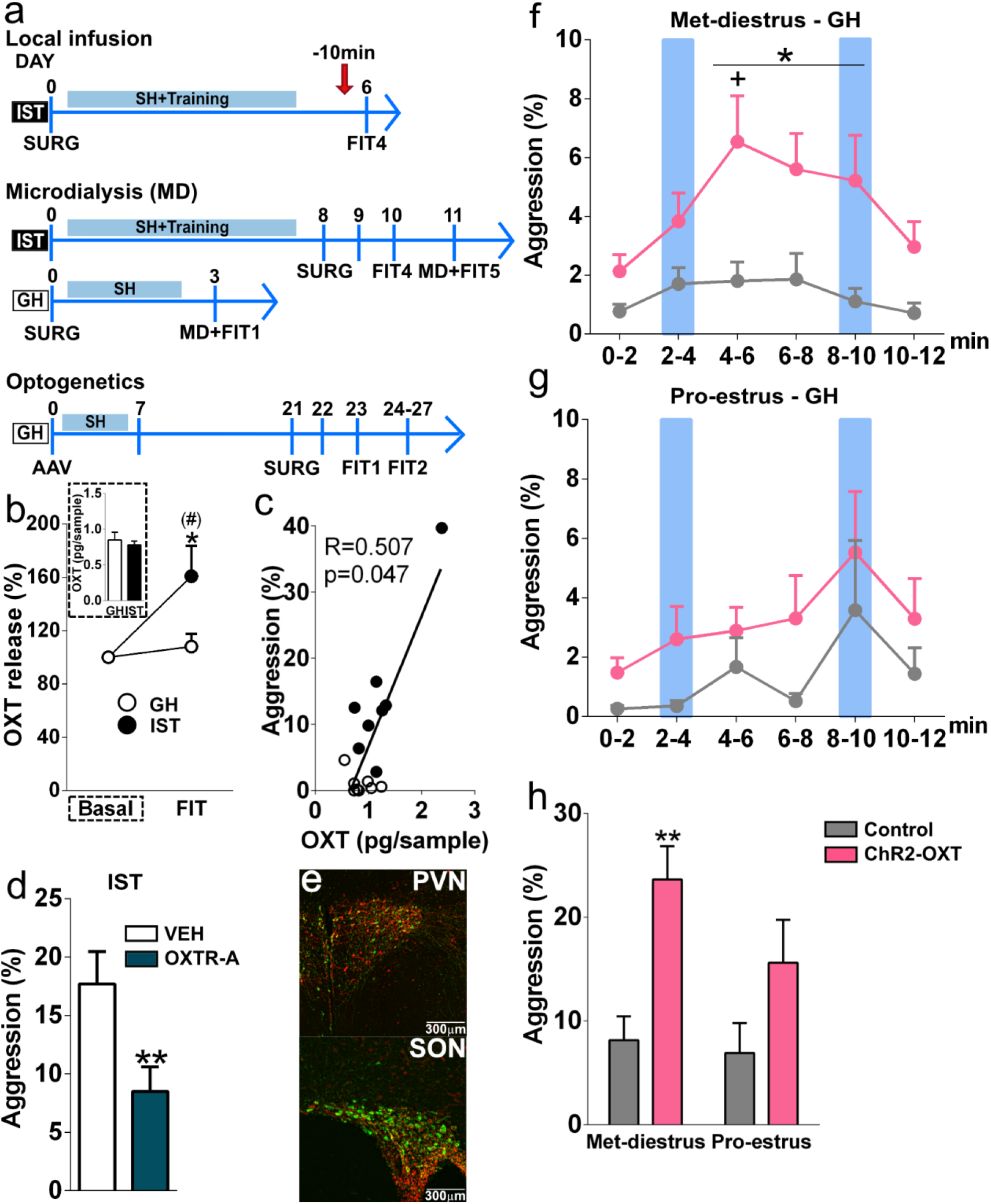
The pro-aggressive effect of OXT is mediated in the vLS. **a** Scheme illustrating the experimental design for pharmacological, microdialysis and optogenetic experiments targeting the OXT system (AAV= adeno-associated virus microinfusion into the paraventricular (PVN) and supraoptic (SON) nuclei; arrow= drug infusions; FIT= female intruder test; GH= group-housed; IST= isolated and trained; MD= microdialysis; SH= single housing; SURG= surgery; OXT= oxytocin; OXTR-A= OXT receptor antagonist; vLS: ventral part of the lateral septum). **b** IST, but not GH females showed an increased rise in OXT release in the vLS during the FIT indicated by increased OXT content in 30-min microdialysates (One sample Student’s t-test IST: t_(7)_=2.65, p=0.033; GH: t_(7)_=0.83, p=0.43), thus OXT release during FIT tended to be higher in IST compared with GH rats (t_(14)_=2.12, p=0.053). Insert shows that absolute OXT content in microdialysates sampled under basal conditions did not differ between the groups (t_(14)_= 0.54, p=0.60). **c** Aggression displayed during the FIT correlated with OXT release in the vLS (Spearman’s correlation r=0.507, p=0.047). **d** OXTR-A (100ng/0.5μl) infusion into the vLS reduced total aggression (two-tailed Student’s t-test t_(26)_=2.58, p=0.016) in IST females. **e-h** Optogenetic stimulation (indicated by blue columns) of OXT axons in the vLS of GH females during the FIT. **e** Confirmation of virus infection in OXT neurons of the PVN (up) and SON (down). OXT-neurophysin I staining: green; mCherry (virus): red. Scale bars 300μm. Blue-light stimulation of channelrhodopsin (ChR2)-OXT fibers enhanced aggressive behavior in **f** met-diestrus (two-way ANOVA followed by Bonferroni factor: time: F_(5,75)_=2.72, p=0.026; virus: F_(5,75)_=20.03, p=0.0004; timexvirus: F_(5,75)_=1.056, p=0.392), but not in **g** pro-estrus females (tim4: F_(5,70)_=2.84, p=0.02; factor virus: F_(1,14)_=2.73, p=0.12; virus x time: F_(5,70)_=0.02, p=0.97). **h** Cumulative analyses show that light stimulation enhanced aggression only in ChR2-OXT females in the met-diestrus phase of the cycle (factor virus: F_(1,15)_=13.06, p=0.0026; estrus-cycle: F_(1,15)_= 2.07, p=0.17; virus x estrus-cycle: F_(1,15)_= 1.114, p=0.31). Data are shown as mean+SEM. ^(#)^p=0.05 vs GH; ^*^p<0.05, ^**^p<0.01 vs either vehicle, baseline or ChR2 control; ^+^p<0.05 vs 0-2 time-point. Microdialysis: n=8; OXTR-A: n=13-15; Optogenetics: n= 8-9.

*In vivo* monitoring of OXT release within the vLS revealed an increased release in IST, but not GH rats during FIT exposure, which was found to correlate with the total aggression displayed (Figure 4b-c, Supplementary Table 2-3), resulting in higher levels of local OXT release observed during the FIT in IST females compared with GH controls. In order to test for the involvement of locally released OXT and subsequent OXTR-mediated signaling in the vLS in female aggression, we bilaterally infused an OXTR-A into the vLS of IST rats, 10 min prior to the FIT. Blockade of vLS OXTR resulted in decreased total aggression (Figure 4d), threat and offensive grooming (Supplementary figure 5).

Further evidence for the involvement of septal OXT neurotransmission in promoting female aggression was provided by optogenetic stimulation of oxytocinergic axons and, thereby, local OXT release, in the vLS of GH rats. To this end, GH rats were infected with a channelrhodopsin (ChR2) rAAV in the PVN and SON, which is expressed under the control of an OXT promoter fragment (Figure 4e). Similarly to the GqDREADD rAAV, the ChR2 rAAV showed a high degree of cell-type specificity. 64.4% of cells in the PVN and 75.1% cells in the SON were positive for both OXT and mCherry. The specificity of the virus for targeting OXT neurons has been proven before^40^, although in the present experiment some limited number of mCherry-positive cells (10.4±1.6 cells/per section, 176 cells in total) outside the PVN and SON devoid OXT immunosignals.

In agreement with the pro-aggressive effects of *endogenous* OXT (Figures 2a-b and 3), blue-light stimulation of vLS OXT axons during the FIT increased the level of aggression in a time- and estrus-cycle-dependent manner (see Figure 4f-h). Cumulative analyses showed a main effect of the viral infection, i.e. only metestrus/diestrus, but not proestrus/estrus, females that expressed ChR2 and received blue-light stimulation, thereby releasing OXT in the vLS, exhibited higher aggression compared to controls (Figure 4h, Supplementary movie 2 and 3 and figure 4).

### AVP exerts an anti-aggressive effect within the dLS

To localize the anti-aggressive effects of AVP identified above, we used intracerebral microdialysis and neuropharmacology within the dLS (Figure 5a), as local V1aR binding was decreased in highly aggressive IST females and negatively correlated with aggression levels (Figure 2h-i).

**Figure 5.**
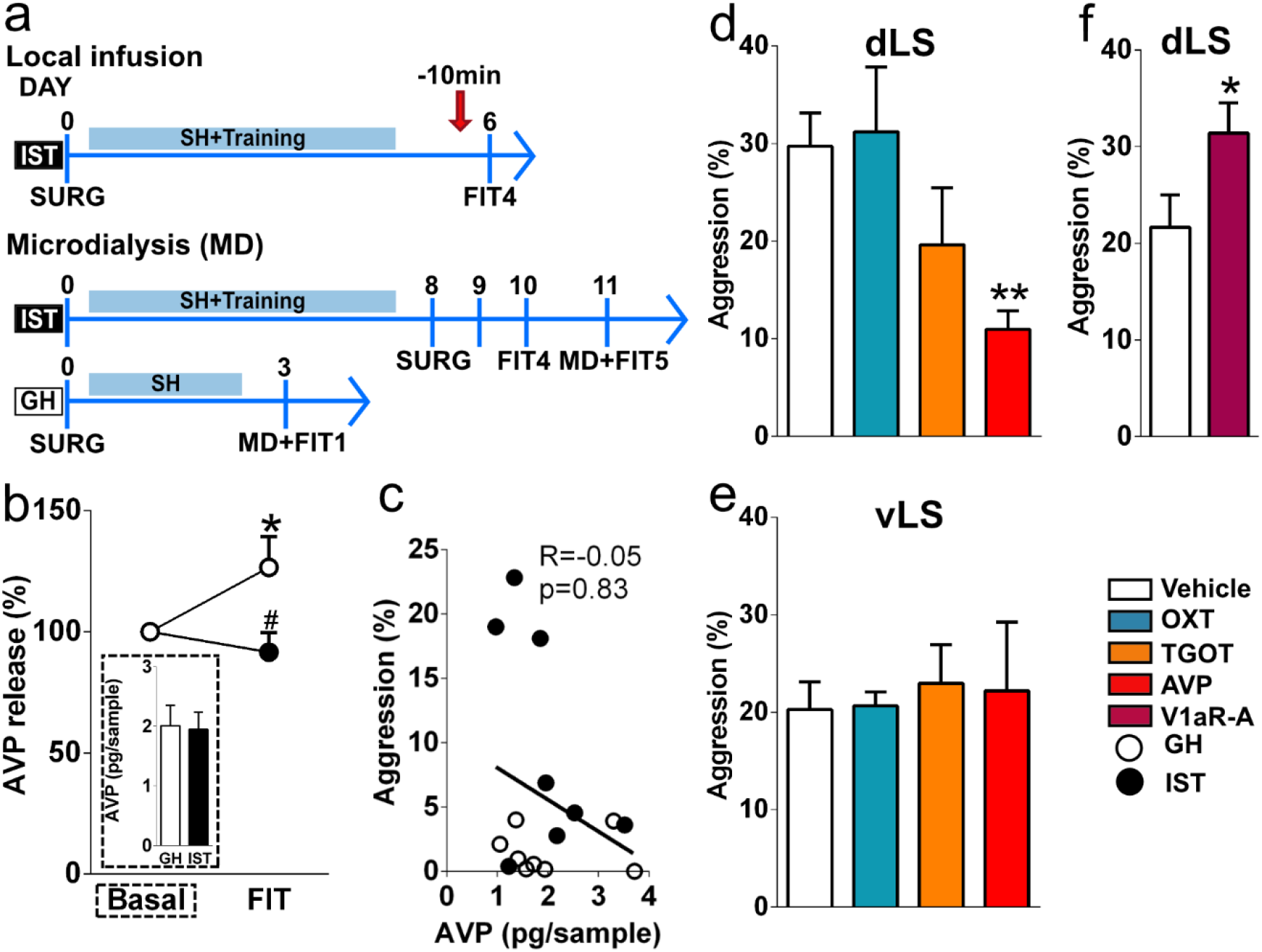
AVP exerts anti-aggressive effects within the dLS. **a** Scheme illustrating the experimental design for pharmacology and microdialysis experiments targeting the AVP system (AVP= vasopressin; arrow= drug infusions; FIT= female intruder test; GH= group-housed; i.c.v.= intracerebroventricular; IST= isolated and trained; MD= microdialysis; SH= single housing; SURG= surgery; dLS: dorsal part of the lateral septum; OXT= oxytocin; TGOT= [Thr4,Gly]OXT, OXT receptor agonist; vLS: ventral part of the lateral septum; V1aR-A= V1aR receptor antagonist). **b** GH, but not IST females showed an increased rise in AVP release in the dLS during the FIT indicated by increased AVP content in 30-min microdialysates (Wilcoxon Signed Rank test GH: W_(9)_=45, p=0.0039; IST: W_(7)_=−4.00, p=0.81), thus AVP release during the FIT was higher in GH than in IST rats (Mann-Whitney U-test U=6.00, p=0.0052). Insert shows that absolute AVP content in microdialysates sampled under basal conditions did not differ between the groups. **c** Female aggression did not correlate with AVP content in the dLS during the FIT (Pearson’s correlation r=−0.197, p=0.46). **d** Infusion of AVP, but not OXT or TGOT (all at: 0.1 ng/0.5 μl), into the dLS decreased total aggression in IST females (F_(3,30)_= 7.292, p=0.0008). **e** AVP infusion into the vLS did not change aggression in IST females (F_(3,25)_=0.11, p=0.95). **f** Local blockade of V1aR (100ng/0.5μl) prior to the FIT increased female aggression in IST rats (two-tailed Student’s t-test t_(20)_=2.14, p=0.045). Data are shown as mean+SEM. ^#^p<0.05 vs GH; ^*^p<0.05, ^**^p<0.01 vs either vehicle or baseline. Microdialysis: n= 8-9; AVP, OXT, TGOT dLS: 6-12 vLS: 6-8; V1aR-A: n=11-12.

AVP release was monitored within the dLS of GH and IST rats under basal conditions and during exposure to the FIT. Whereas in GH rats, AVP release was found to significantly increase to 130 %, such a rise was not found in IST females resulting in lower AVP content during the FIT compared with GH controls (Figure 5b, Supplementary Table 2-3). However, no correlation between local AVP release and aggressive behavior was identified (Figure 5c). Next, to prove that AVP acts on V1aRs specifically in the dLS to inhibit female aggression, we infused AVP, TGOT (a selective OXTR agonist), or OXT either into the dLS, where we identified predominantly V1aRs, or into the vLS, where predominantly OXTRs were found (Figure 2e and j). Only bilateral infusions of AVP, into the dLS of IST rats resulted in decreased total aggression, keep down, and offensive grooming (Figure 5d and Supplementary figure 5). In contrast, none of the treatments in the vLS affected aggression (Figure 5e and Supplementary Table 2). Surprisingly, V1aR-A administration into the dLS alone had a moderate effect on enhancing aggression in IST rats (Figure 5f and Supplementary Figure 5).

### Spontaneous activity in neurons in the dLS and vLS is differentially modulated by activation of OXTRs

The LS consists mostly of GABAergic neurons grouped into different subnuclei according to the expression of different neuropeptides and receptors, most importantly V1aRs and OXTRs^34,41^. It has been reported that spontaneous activity in these neurons is modulated by AVP in multiple ways^42^. In our behavioral paradigm, activation of OXTRs in the vLS enhanced, whereas activation of V1aRs in the dLS rather reduced female aggression. To test whether such subregion- and neuropeptide-specific effects can also be observed at the level of network activity, we recorded spontaneous activity of GABAergic dLS and vLS neurons in brain slices from juvenile Venus-VGAT rats (whole-cell voltage-clamp at −60mV, biocytin-labelling) and investigated OXTR- and V1aR-mediated effects by applying either TGOT (1μM) or a combination of OXTR-A (10μM) and AVP (1μM) to the bath. We also characterized these two neuronal populations with respect to morphological and molecular parameters (Figure 6a).

**Figure 6.**
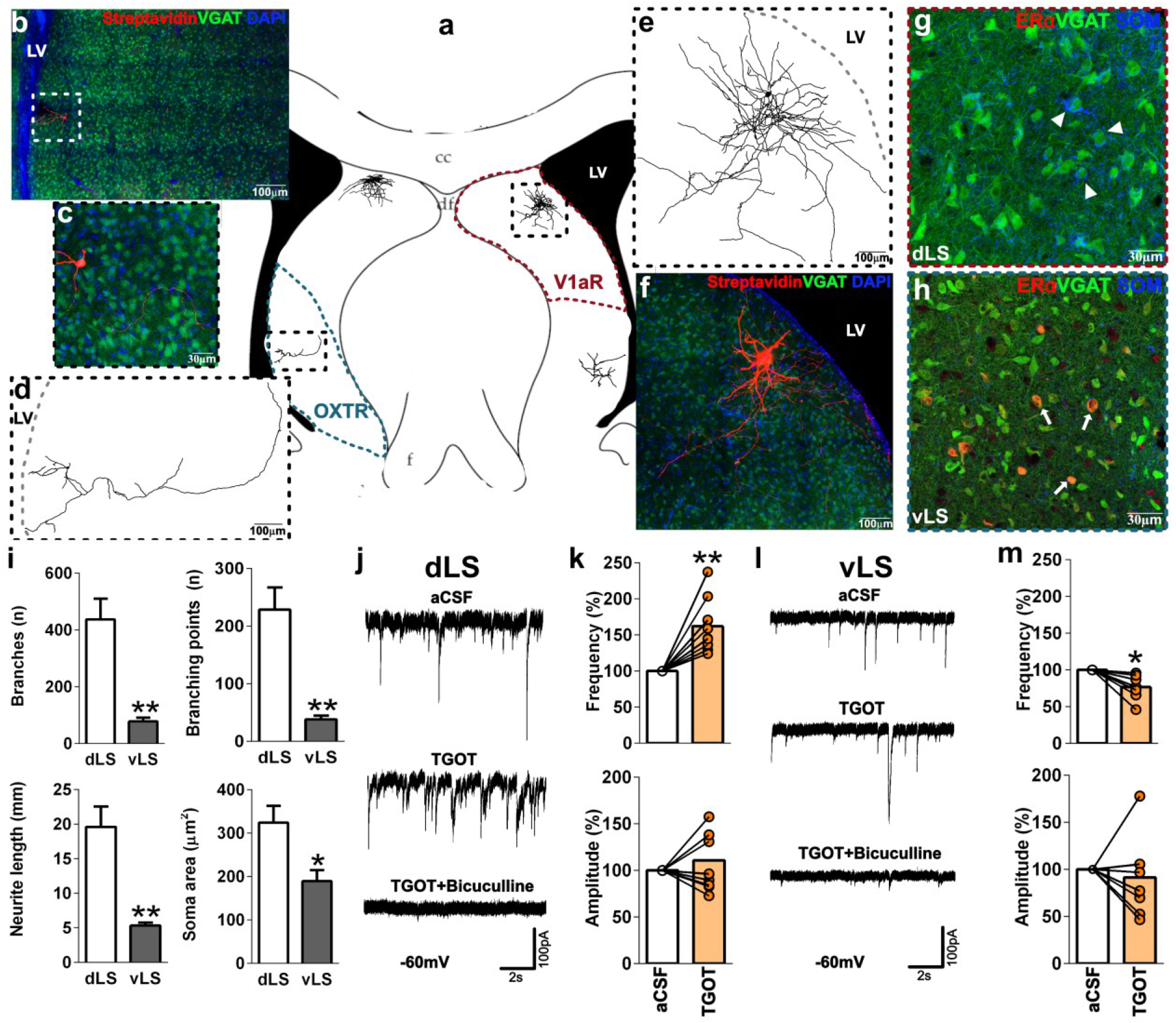
Spontaneous activity in neurons of the dLS and vLS is differentially modulated by activation of OXTRs. **a** Scheme indicating the receptor binding-specific delimitation of the vLS (OXTR) and dLS (V1aR) used during voltage-clamp experiments, including two representative dLS and vLS cells. **b** Maximal z-projection of biocytin-filled vLS cell (streptavidin-CF633) overlaid with a single z-plane of VGAT and DAPI. **c** Magnification of cell body from the cell shown in **b**. **d** Neurite reconstruction of cell shown in **b**, and **e** of the cell shown in **f**; the gray-dotted line indicates the border of the lateral ventricle (LV). **f** Maximal z-projection of a biocytin-filled dLS (streptavidin-CF633) cell overlaid with a single z-plane of VGAT and DAPI. **g** Magnification of a single z-plane indicating the presence of somatostatin (SOM) cell bodies (arrowheads) only in the dLS. **h** Magnification of a single z-plane indicating the presence of ERα-positive cell bodies (arrows) only in the vLS. **i** Morphological characterization of LS neurons. dLS cells exhibited more neurite branches (two-tailed Student’s t-test t_(4)_= 4.84, p=0.0072) and branching points (t_(4)_= 4.85, p=0.0072) longer neurite length (t_(4)_= 4.80, p=0.0078), and wider soma areas (Mann-Whitney U-test U= 1.0, p=0.016) than vLS cells. Representative spontaneous current traces during TGOT (1μM) and bicuculline (50μM) bath application in dLS **j** and vLS **l** cells. **k** TGOT increased sIPSC frequencies in dLS cells (Wilcoxon signed rank test W_(9)_=55, p=0.002) and **m** decreased sIPSC frequencies in vLS cells (W_(7)_=−36, p=0.0078). TGOT had no effect whatsoever on the sIPSC amplitude independent of the subregion. Data are shown as mean+SEM. ^*^p<0.05, ^**^p<0.01, vs either dLS or aCSF. dLS n=10; vLS n=8.

Neurons in the dLS exhibited a higher morphological complexity compared to vLS neurons (Figure 6a-f), as indicated by higher numbers of neurite branches and branching points (Figure 6i). Apart from the specific expression of V1aRs and OXTRs, dorsal and ventral LS neurons also differed regarding the expression of other markers: somatostatin-positive cell bodies were only found in the dLS (Figure 6g), whereas estrogen receptor α (ERα) expressing cells were exclusively located in the vLS (Figure 6h), further reinforcing that vLS and the dLS neurons are distinct populations.

While excitatory spontaneous activity was not observed under our recording conditions, in line with previous observations^42^, selective activation of OXTRs in the vLS differentially affected the spontaneous inhibitory activity in dLS versus vLS (Figure 6j and l). In the dLS cells, the selective OXTR agonist TGOT caused an increased frequency of spontaneous IPSCs (sIPSCs). sIPSCs were entirely blocked by further addition of bicuculline (50μM, selective GABA-A antagonist) (Figure 6j-k), which confirmed the exclusive GABAergic origin of these currents. In agreement with those results, blockade of OXTRs by selective OXTR-A in the dLS decreased sIPSC frequency (Supplementary figure 6a-b). Conversely, in the vLS neurons responded to TGOT with a decreased sIPSC frequency (Figure 6l-m) indicating a reduced inhibitory tone following OXTR activation. Again, sIPSCs were entirely blocked by bicuculline.

V1aR activation increased sIPSC frequency specifically in the dLS neurons, whereas no consistent effect was seen in vLS cells (Supplementary figure 6).

Altogether, our data demonstrate that activation of OXTRs in the vLS concomitantly weakens the tonic inhibition of vLS GABAergic neurons and strengthens tonic inhibition of dLS GABAergic neurons. We also could confirm subregion-specific effects of AVP, since only dLS neurons were affected by V1aR activation, in line with behavioral experiments where AVP effects were also observed only in the dLS. In addition, dLS neurons showed extended dendritic arborizations and other cell markers than vLS neurons.

### An intrinsic GABAergic circuit within the LS regulates female aggression

After we have shown that i) activation of vLS OXTR increases, whereas dLS V1aR decreases aggression, and ii) spontaneous inhibition is increased in dorsal and decreased in ventral cells by OXTR activation, we hypothesized that those subregions would also be differentially activated after an aggressive encounter. Therefore, we compared the neuronal activity within the vLS and dLS of Venus-VGAT GH and IST rats after exposure to the FIT using pERK as a neuronal activity marker^12,14,15^.

We found striking regional differences in neuronal activity in the LS of highly aggressive IST females compared to low aggressive GH rats (Figure 7a-d). In detail, in the dLS FIT exposure and the display of high aggression by IST rats resulted in a decreased total number of both pERK-positive and VGAT/pERK-positive cells (Figure 7a-b and e), with a negative correlation found between the amount of double-labeled cells in the dLS and aggression (Figure 7f). In contrast, in the vLS of IST females, we found a trend towards more pERK-positive cells and an increased number of VGAT/pERK positive cells (Figure 7c-d and g), which did not correlate with aggression (Figure 7h and Supplementary Table 2-3).

**Figure 7.**
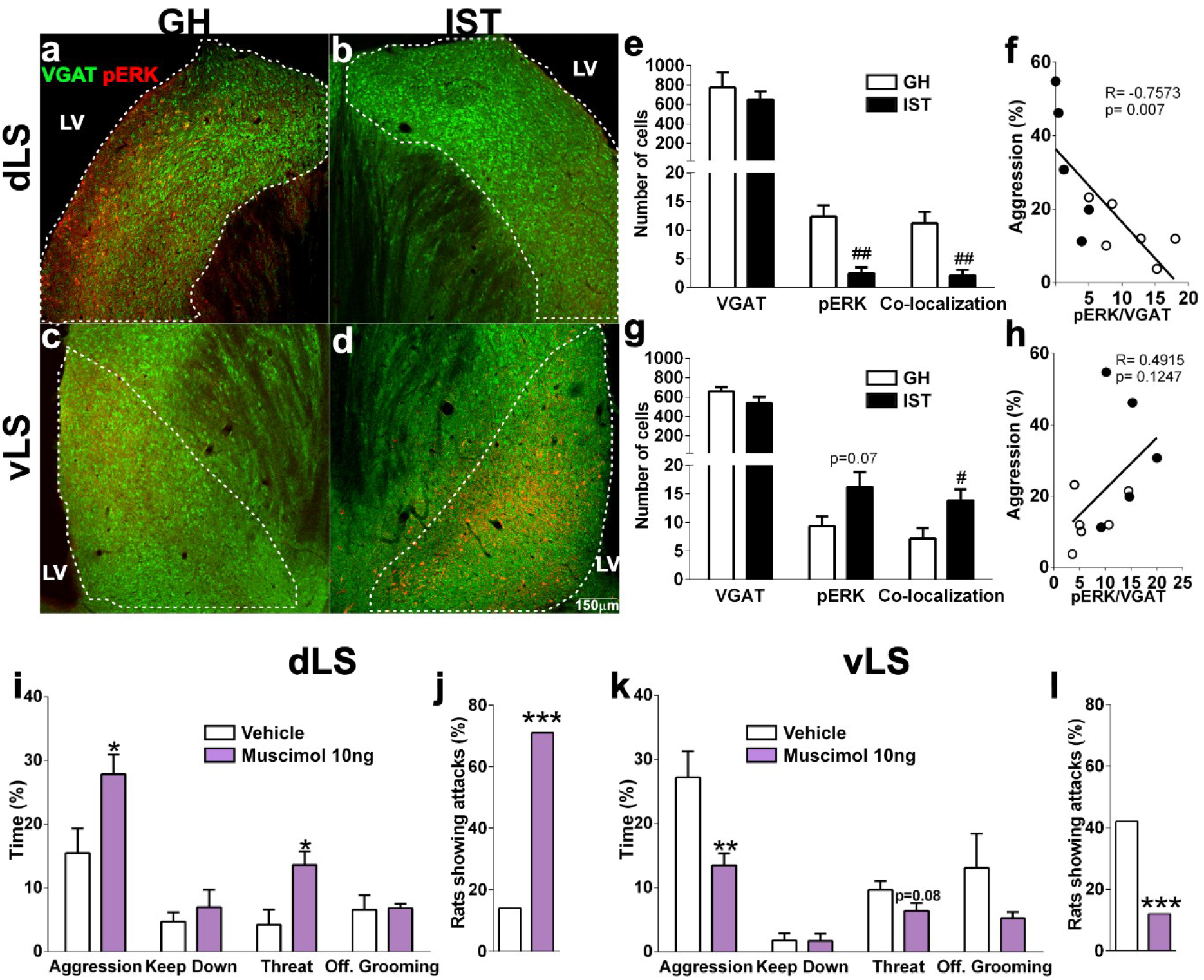
An intrinsic GABAergic circuit within the subregions of the LS regulates female aggression. **a-d** Example average z-projects showing pERK (Alexa594) immunostaining in Venus-VGAT females after exposure to the female intruder test (FIT). **a** dorsal LS (dLS) of a group-housed (GH), i.e. low aggressive female, **b** dLS of an isolated and trained (IST), i.e. high aggressive female, **c** ventral LS ( vLS) of a GH female, **d** vLS of an IST female (LV=lateral ventricle; VGAT= vesicular GABA transporter;). **e-h** Neuronal activity in the LS reflected by pERK staining after FIT exposure: **e** In the dLS of IST females less pERK-positive cells (two-tailed t-Student’s test t_(9)_=4.20, p=0.0023) and fewer pERK/VGAT co-localized cells were found (t_(9)_=3.75, p=0.004). **f** Aggression negatively correlated with the number of pERK/VGAT-positive cells in the dLS (Pearson’s correlation r=−0.746, p=0.008). **g** In the vLS of IST females a tendency of more pERK-positive cells (t_(9)_=2.07, p=0.068) and a higher number of pERK/VGAT-positive cells was found (T_(9)_=2.28, p=0.049). **h** Aggression did not correlate with the number of pERK/VGAT-positive cells in the vLS (r=0.4915, p=0.1247). **i** Infusion of muscimol (10ng/0.5μl) into the dLS increased total aggression (t_(12)_= 2.52, p=0.027) and threat (Mann-Whitney U-test U=5.00, p=0.011) in GH rats. **j** Inhibition of the dLS also enhanced the percentage of rats showing attacks (Fisher exact test, p<0.0001). **k** Muscimol in the vLS decreased total aggression (t_(13)_=3.191, p=0.0071) and tended to decrease threat behavior (t_(13)_= 1.832, p=0.090). **l** Inhibition of the vLS also reduced the percentage of rats showing attacks (p<0.0001). Data are shown as mean+SEM. ^#^p<0.05, ^##^p<0.01 vs GH; ^*^p<0.05, ^**^p<0.01, ^***^p<0.0001 vs vehicle. Neural activity: n=5-6; Muscimol: dLS: n=7 vLS: n=7-8.

To further confirm the dLS as an anti-aggressive center versus the vLS as a pro-aggressive center, and consequently to create a causal link between neuronal activity and behavior, we infused rats with a selective GABA-A agonist muscimol 10 min before the FIT. As predicted, inactivation of the dLS in GH rats enhanced aggression, threat behavior and the percentage of rats showing attacks (Figure 7i-j and Supplementary Table 2-3). In contrast, opposing effects were seen in the vLS of IST rats: muscimol decreased aggression and the percentage of rats showing attacks. (Figure 7k-l and Supplementary Table 2-3).

## Discussion

Aiming to study neural mechanisms of female aggression, we have established a novel behavioral paradigm combining social isolation and aggression-training, which reliably exacerbated the mild levels of aggression typically displayed by GH female Wistar rats. Additionally, we have shown that female aggression is controlled by a fine-tuned balance between OXT, AVP, and GABA neurotransmission within two distinct neuronal populations of the LS, i.e. the dLS and vLS, with contrasting subregional effects of OXT and AVP (Figure 8). In detail, the display of high aggression by IST females during the FIT was accompanied by elevated OXT release within the vLS, which was also reflected by higher OXT levels in their CSF. Together with the fact that central OXT release correlated with aggression, this indicates the involvement of septal OXT in the high aggression displayed by IST rats, which we further confirmed by complementary pharmacological, chemogenetic and optogenetic approaches: (i) Pharmacological blockade of OXTR either i.c.v. or locally in the vLS attenuated aggression in IST rats, whereas (ii) chemogenetic and optogenetic stimulation of OXT release within the brain, and specifically within the vLS, heightened the aggression level of low aggressive and non-receptive GH females.

**Figure 8.**
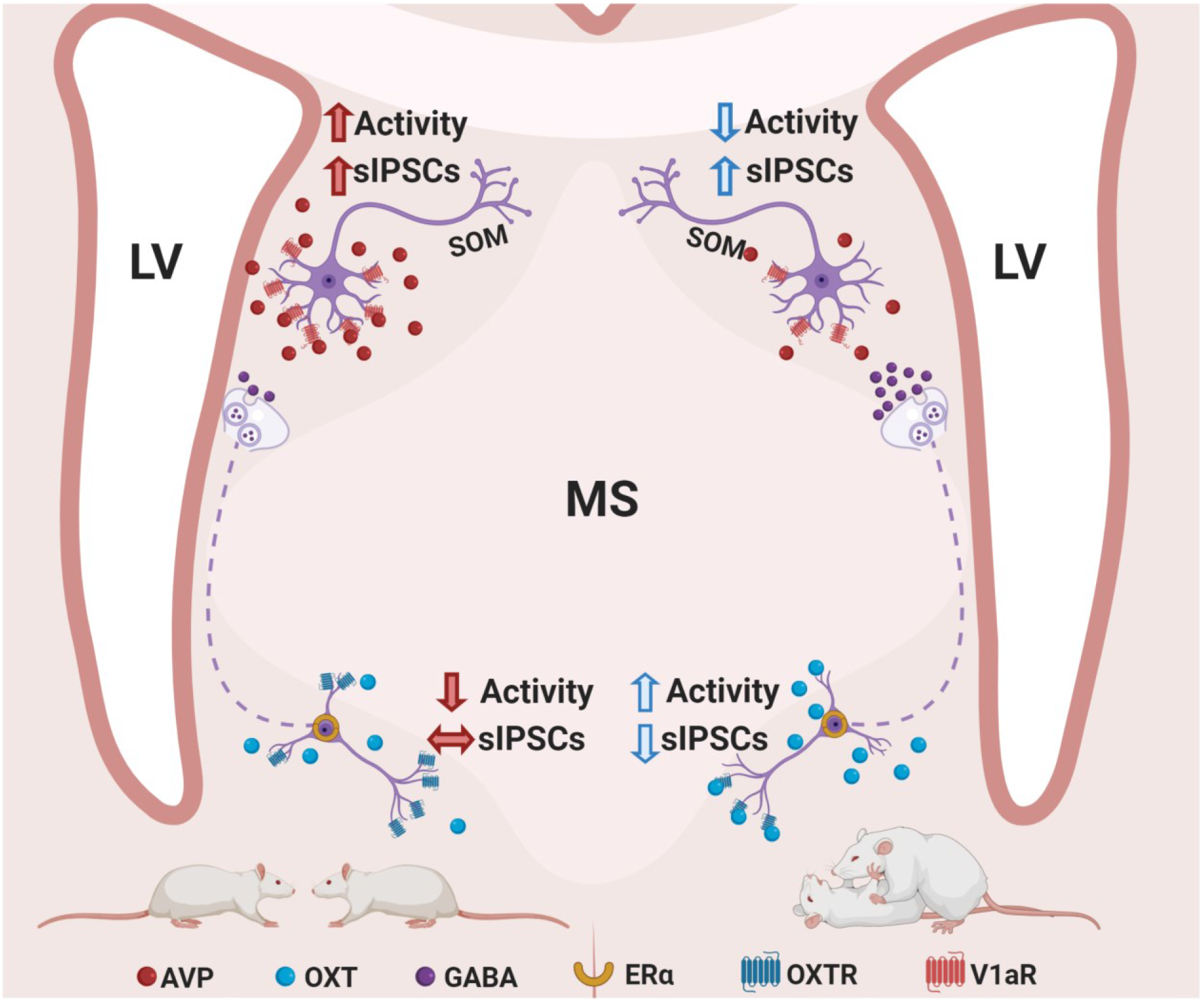
The balance between OXT and AVP regulates the inhibitory tonus within the LS in order to control female aggression in Wistar rats. The scheme depicts the main findings of this manuscript (partially drawn using https://biorender.com). Further, it hypothesizes how alterations in OXT and AVP release may impact a microcircuit in the LS in order to generate aggressive display in female Wistar rats. Dorsal and ventral cells are shown in proportion to their real size (AVP= vasopressin; ERα= estrogen receptor α; IPSC= inhibitory post-synaptic current; LS= lateral septum; LV= lateral ventricle; MS= medial septum; OXT= oxytocin; OXTR= oxytocin receptor; SOM= somatostatin; V1aR= V1a receptor). On the left side, is depicted how, the brain of a GH female responds to an intruder: a combination of low OXT release ventrally and high AVP release dorsally evokes an increased activity of the dLS, thereby reducing aggression. On the right side is depicted how the brain of an IST female responds to an intruder: a combination of high OXT release ventrally and low AVP release dorsally culminates in increased tonic inhibition of dLS (dotted line indicates hypothetical pathway) and increased activation of the vLS, promoting aggression.

Concerning the involvement of the brain AVP system in female aggression, we revealed a completely different picture. Highly aggressive IST females showed blunted AVP release within the dLS during an aggressive encounter, which was also reflected by lower AVP content in the CSF. As further proof of an inhibitory effect of AVP on female aggression, infusion of AVP either i.c.v. or directly into the dLS reduced the aggression seen in IST rats, whereas blockade of V1aR in the dLS exacerbated their aggression. Furthermore, IST rats exhibited reduced V1aR binding in the dLS, and aggressive behavior negatively correlated with V1aR binding.

OXT and AVP have been repeatedly described as important neuromodulators of social behaviors in rodents, acting either synergistically or antagonistically depending on the social context and sex^8,9,22,24,36,43^. In males, reduced aggression has been reported in rats after i.c.v. infusion with synthetic OXT ^23^ and in mice after optogenetic stimulation of hypothalamic OXT neurons ^43^. In contrast, the few studies on OXT-effects on female aggression rather revealed pro-aggressive or antisocial effects, for example in female rhesus monkeys^44^, non-lactating women^8,45^, and in lactating female rats^18,33^. Although the factors motivating the display of aggression and its severity likely differ in lactating versus virgin females^5,33,46,47^, we should highlight that from an evolutionary point of view, co-opting the same neuropeptidergic systems for promoting aggression in lactating females to protect the offspring, and in virgin females to protect their territory, or to get access to resources, makes sense; especially knowing that high activity of the brain OXT system reflected by elevated neuropeptide synthesis, release and binding^18,24,33,36^ is known to underlie maternal aggression^24,33,36,48^. Further evidence for a sex-specific effect of OXT on social behavior comes from studies on social motivation, where OXT is essential for naturally occurring social preference behavior in male rats and mice^49^, but not in virgin female rats^50^. Thus, our results on a pro-aggressive effect of OXT in virgin females add an important piece of evidence to the sex-dimorphic effects of OXT on social behaviors.

In our experiments, female aggression was found to be dependent upon the estrus cycle. Indeed, a previous study has shown low aggression in estrus rats^51^, as we have seen in the IS proestrus/estrus females. Additionally, chemo- and optogenetic stimulation of OXT release either centrally or in the vLS elevated aggression levels exclusively in metestrus/diestrus GH rats. In this context, it is of note that OXTR expression and binding in regions involved in the regulation of aggression, such as the VMH, bed nucleus of *stria terminalis* (BNST), and LS, undergo dynamic changes during the estrus cycle in a sex steroid-dependent manner^36,52^. Furthermore, sex steroids affect OXT expression and release as shown *in vitro*^53^ and *in vivo*^54^. Although one could hypothesize that activation of local ERα in the vLS is counteracting the effects of OXT in females, the extent of how the estrus cycle is involved in the modulation of the vLS neurons and, consequently, on local OXT and AVP actions on female aggression remains to be studied.

While several pieces of evidence support pro-aggressive effects of AVP in males^9,22,55,56^, conflicting data supporting an anti-aggressive effect of AVP have been described in animal models of high and abnormal aggression. For example, male rats bred for low anxiety and characterized by excessive and abnormal aggression^31^ show low levels of AVP release in the LS during the resident-intruder test^21^. Also, aggressive male mice with a short attack latency exhibit decreased AVP innervation of the LS^57^, thereby demonstrating not only an association between abnormal and high aggression, but furthermore a blunted activity of the septal AVP system. In agreement, high aggression displayed by dominant males has also been linked to reduced septal AVP signaling, i.e. alpha male mice showed decreased V1aR binding in the LS, when compared to subordinate males^32^, and synthetic AVP was able to flatten dominant behavior in rhesus macaques^58^. In lactating females, the link between AVP and aggression seems to be complex as well. Whereas *synthetic* or *endogenous* AVP has been shown to decrease maternal aggression in Sprague-Dawley rats^19,20^, contrasting and probably anxiety-dependent effects have been described in the central amygdala of high anxiety Wistar rats, where AVP release was associated with maternal aggression^17^. In accordance with our data, serenic effects of AVP have also been shown in non-lactating hamsters^9,59^ and rhesus macaques^44^. Interestingly, recent studies in humans have highlighted the pro-social role of AVP as a potential treatment for social dysfunctions, since intranasal AVP was able to increase risky cooperative behavior in men^60^ and to enhance social skills in autistic children^61^.

After identification of the specific pro-aggressive effect of OXT in the vLS, and anti-aggressive effect of AVP in the dLS of female Wistar rats (Figure 8), in the final set of experiments we were able to link these neuropeptidergic actions with specific regional effects in neuronal activity after an aggressive encounter. In detail, elevated pERK expression reflecting high neuronal activity was found in the vLS of highly aggressive IST females, whereas pERK expression was elevated in the dLS of low aggressive GH rats. These findings were functionally validated by manipulation of local GABAergic neurotransmission using local muscimol infusions, demonstrating that dLS neurons inhibit, whereas vLS neurons seem to promote female aggression. Subnetwork-dependent responses were also seen after OXTR and V1aR activation *in vitro*. TGOT decreased the inhibitory spontaneous activity ventrally, whereas it increased it dorsally. Thus activation of OXTRs located exclusively in the vLS leads to enhanced GABAergic inhibition of GABAergic neurons in the dLS, possibly facilitating aggression. Additionally, AVP increased sIPSCs frequency dorsally, but did not affect sIPSCs ventrally. This observation refines the results from an earlier study^42^ showing that V1aR activation indirectly enhances the inhibitory neurotransmission of a majority of LS neurons, notwithstanding the dorso-ventral localization of the recorded cells. Thus, according to our results, both anti-aggressive and synaptic effects of AVP appear to be mediated exclusively within the dLS.

The role of the LS in suppressing aggression has been described for decades^10,25–30^. However, only one recent paper has shown the existence of local microcircuits within the LS regulating aggression in male mice^56^. Supporting the pro-aggressive role of AVP in males^9,22,24,55^, V1b receptor activation on presynaptic terminals of hippocampal fibers to the LS increased aggression via stimulation of inhibitory interneurons in the dLS projecting to the vLS. Although this data contrasts with ours, we have to keep in mind that i) the regulation of aggression within the LS, via OXTRs and V1aRs, might be underlined by sex- and species-dependent mechanisms, as males and females^34^, as well as mice and rats^32,34^, differ in their receptor binding, (ii) we investigated the role of V1aRs, but not V1bRs and differential effects on aggression are likely, and iii) the involvement of endogenous AVP acting on V1bRs has not been shown^56^. Based on our finding that V1aR activation increases sIPSCs frequency exclusively in the dLS, we could hypothesize that V1aR activation might overrule V1bR effects by inhibiting V1bR-responsive interneurons, thereby decreasing aggression. Accordingly, in our female aggression paradigm, activation of V1aRs in the dLS seems to overshadow any possible pro-aggressive V1bR-mediated action, since AVP administration in the dLS was able to mimic the anti-aggressive effects of i.c.v. AVP in IST females. Indeed, AVP has been shown to present a higher affinity to V1aRs over V1bRs *in vitro*^37^. Nevertheless, future studies should address how the interplay between V1aR and V1bR activation within the dLS affects female aggression.

Supporting our findings, a neurocircuit involving contrasting effects of OXTR and V1aR activation has been described in the context of fear in the central amygdala, wherein activation of putatively OXTR-positive cells led to inhibition of V1aR-positive cells via GABAergic transmission, which abolished fear-related behavioral effects of V1aR activation^40,62^. To the best of our knowledge, such a mechanism has never been described before in the context of aggression. In further support of our finding that high levels of OXT in the vLS leads to increased tonic inhibition of the dLS via GABAergic projections, neural projections from the vLS towards the dLS have also been described in male Wistar rats^63^. Additionally, blockade of GABA-A receptors in the dLS of lactating mice reduced maternal aggression^30^. Thus, aggression in animals with a higher OXT system activity ^33,36^ is at least partially modulated by increased GABA neurotransmission in the dLS.

Taken together, our results are among the first to shed light on the neurobiological mechanisms underlying female aggression. We have shown that the level of aggression expressed by virgin female Wistar rats is determined by a fine-tuned balance between OXT- and AVP-mediated neuromodulation in septal sub-networks. Disturbances in this balance, such as a shift towards a more dominant OXT or AVP neurotransmission, lead to distinct aggression responses. Dorsal V1aR-expressing neurons of the LS seem to be pivotal for the AVP-induced inhibition of aggression whereas ventral neurons in the OXTR-expressing LS appear to be the main generators of OXT-induced aggression. Additionally, we propose a mechanism by which activation of OXTRs reduces tonic inhibition of vLS neurons, which in turn may enhance their GABA transmission onto the dorsal cells and thus ultimately may reduce septal gating on aggression (Figure 8).

## Methods

### Ratlines and animal care

Adult female Wistar rats (10-14 weeks old) bred in the animal facilities of the University of Regensburg (Germany) were used for behavioral experiments. Intruders, female Wistar rats were obtained from Charles Rivers Laboratories (Sulzfeld, Germany) and kept in groups of 3 to 5 animals in a separate animal room. All rats were kept under controlled laboratory conditions (12:12 h light/dark cycle; lights off at 11:00, 21±1°C, 60±5% humidity) with access to standard rat nutrition (RM/H, Sniff Spezialdiäten GmbH, Soest, Germany) and water *ad libitum*. For patch-clamp and pERK immunohistochemistry analyses, Venus-VGAT rats (VGAT: vesicular GABA transporter; lineVenus-B, W-Tg(Slc32a1-YFP*)1Yyan)^64^ that were bred in the animal facilities of the University of Regensburg were used. All procedures were conducted following the Guidelines for the Care and Use of Laboratory Animals of the Local Government of Oberpfalz and Unterfranken.

### Female Intruder test (FIT)

The FIT was performed in the early dark phase (between 12:00-16:00) under dim red light conditions. An unfamiliar female intruder weighing between 10-20% less than the resident^12^ was released into the home cage of the resident for 10 minutes. The test was videotaped for later analysis by an experienced observer blind to treatment using JWatcher event recorder Program^65^. The percentage of time the resident spent with four main behavioral aspects was scored: i) aggressive behaviors, i.e., keep down, threat behavior, offensive grooming, and attacks; ii) neutral behaviors, i.e., exploring and investigating the home cage, drinking, eating and self-grooming; iii) social behaviors, i.e. non-aggressive social interactions, sniffing, following; and iv) defensive behavior, i.e. submissive posture, kicking a pursuing intruder with hind limb. We also monitored sexual behavior (lordosis, hopping, darting and mounting) and immobility. In addition, we scored the frequency of attacks as well as the latency to the first attack. Vaginal smears were taken after the FIT to verify the estrus cycle; all phases of the estrus cycle were included in the study (for detailed behavioral analyses of all experiments please see Supplementary tables 2-3).

### Overview of the *in vivo* experiments

#### Animal groups for OXT/AVP measurements in CSF, plasma, and receptor autoradiographic analyses

Female Wistar rats were split into three different housing conditions: Group-housed (GH) females were kept in groups of 3 to 5 animals per cage, isolated (IS) females were singly housed for 8 days, and isolated-trained (IST) females were also kept singly for 8 days, but from day 5 on they underwent 3 consecutive FITs for aggression-training. On day 9, GH females were singly housed 4 hours prior to the behavioral experiments. One hour after lights went off, GH, IS and IST rats were either exposed to the FIT, whereas control rats were left undisturbed in their home cage. Immediately after the FIT, rats were deeply anesthetized using intraperitoneal (i.p.) urethane (25%, 1.2 ml/kg) to allow CSF collection via puncture of the *Cisterna cerebromedullaris*. After decapitation, brain and trunk blood were collected for receptor binding and hormonal measurements, respectively (Figure 1a).

#### Neuropharmacology design

For pharmacological experiments, female rats were split into GH and IST conditions. All females underwent surgery for intracerebroventricular (i.c.v.) or local cannula implantation. IST females were left undisturbed for recovery for 3 (i.c.v.) or 5 (local) days. The aggression training was performed as described above, except for the fact that residents received a sham-infusion before the FIT to get used either to the i.c.v. or local infusion procedure. GH animals were kept single-housed overnight for recovery and brought back to their original groups the next day until the start of the experiments when they were transferred into an observation cage and single-housed for 4h before the FIT. All animals were handled daily to get used to the infusion procedure. Typically, a cross-over, within-subjects design was used for all i.c.v. agonist experiments, whereas a between-subjects design was used for the local infusions and i.c.v. antagonists experiments. Additionally, to guarantee that all groups had similar average levels of aggression before pharmacological manipulations, we allocated IST rats into the treatment groups according to their average levels of aggression expressed in the third session of training.

#### Microdialysis

OXT and AVP release within the LS was monitored in GH and IST females before and during FIT exposure. After 4 days of training, IST rats had their microdialysis probes implanted into the LS. After one day of recovery, IST subjects received the 5^th^ training FIT to confirm their previous levels of aggression. On the following day, both GH and IST animals underwent the microdialysis procedure. Briefly, rats were connected to a syringe mounted onto a microinfusion pump via polyethylene tubing and were perfused with sterile Ringer’s solution (3.3 μl/min, pH 7.4) for 2 hours before sampling of microdialysates to establish an equilibrium between inside and outside of the microdialysis membrane. One hour after lights went off, three consecutive 30-min dialysates were collected (baseline samples 1 and 2 represented in Figure 4 and 5 as an average of both time-points) and during an ongoing FIT (sample 3). Dialysates were collected into Eppendorf tubes containing 10 μl of 0.1 M HCl, were immediately frozen on dry ice, and stored at −20°C until subsequent quantification of AVP and OXT by radioimmunoassay (for behavioral analyses see Supplementary tables 2-3).

#### Chemo- and optogenetics design

Chemo- and optogenetic experiments targeting the OXT system were only performed in GH rats. After stereotaxic virus infusion into the PVN and SON, animals were kept single-housed for one week to recover. Thereafter, they were again group-housed for two weeks until either the experiment took place (chemogenetic) or they had their optical fiber implanted (optogenetic). Similarly to the pharmacologal experiments, chemogenetic rats were kept in groups and isolated shortly before the dark phase (see above). Subjects received an i.p. infusion of clozapine-N-oxide dihydrochloride (CNO, 2mg/kg) 45 min before the FIT. They were brought back to their original groups directly after the FIT. In the optogenetic experiments, after optical fiber implantation rats were single-housed for three days for recovery and to avoid damaging the fiber. Similarly to the microdialysis experiments, animals were connected to the optogenetic cables two hours before the experiment to get used to the cables. In this experiment, the FIT lasted 12 minutes. Blue-light stimulation (30ms pulses of 30Hz delivered for 2min; in analogy to Knobloch et al., 2012) was delivered at the 2^nd^ and again 8^th^ minute after the beginning of the FIT. For both, chemo- and optogenetic experiments, animals were tested twice at different phases of their estrus cycle: Once in proestrus/estrus, and once in metestrus/diestrus. All animals were transcardially perfused with paraformaldehyde (PFA 4%) for histological verification of the specificity of viral transfections after the last test.

Although the i.c.v. and local pharmacology experiments showed robust serenic effects of AVP (Figure 3 and 5), we decided to not genetically manipulate the AVP system centrally or locally due to three reasons: i) AVP is one of the main players in brain physiology and stimulation of several cell bodies in different brain regions especially in the hypothalamus could disturb homeostasis. In fact, high doses of i.c.v. AVP are known to elicit barrel rotations in rats^66^. ii) AVP neurons are widespread in several nuclei in the rat brain such as the medial amygdala, the BNST, PVN and SON^67^ which makes the infusion of all targeted neurons challenging in terms of animal welfare. iii) We (data not shown) and others^68^ have found low specificity of the AVP directed virus to infect intra- and extrahypothalamic parvocellular AVP neurons, which are known to project to the LS^67^.

#### Neuronal activation after aggression

To compare neuronal activity patterns in the dLS and vLS of IST and GH rats after exposure to the FIT, we used female Venus-VGAT rats (10-14 weeks old). Immediately after FIT exposure rats were deeply anesthetized with isoflurane (ForeneH, Abbott GmbH & Co. KG, Wiesbaden, Germany), followed by CO_2_, transcardially perfused, and brains were harvested for subsequent immunohistochemistry stainings (behavioral data in Supplementary Tables 2-3).

### Stereotaxic surgery

Rats were anesthetized with isoflurane, injected i.p. with the analgesic Buprenovet (0.05 mg/kg Buprenorphine, Bayer, Germany) and the antibiotic Baytril (10 mg/kg Enrofloxacin, Baytril, Bayer, Germany), and fixed in a stereotaxic frame. I.c.v. guide cannulas (21 G, 12 mm; Injecta GmbH, Germany) and microdialysis probes (self-made, molecular cut-off 18 kDa^22^) were implanted unilaterally, whereas local guide cannulas (25 G, 12 mm; Injecta GmbH, Germany) were implanted bilaterally. All cannulas were implanted 2 mm above the target region to avoid lesion of the target region, whereas microdialysis probes had to be implanted directly into the target region, fixed to the skull with two jeweler’s screws and dental cement (Kallocryl, Speiko-Dr. Speier GmbH, Muenster, Germany). The cannulas were closed by a stainless steel stylet (i.c.v. 25 G, local 27 G).

For viral infection in chemo- and optogenetic experiments, a solution of ketamine (100 mg/kg, Medistar, Germany) and xylazine (10 mg/kg, Medistar, Germany) was applied i.p. as anesthesia. After virus delivery, the skin was sutured. For optogenetic experiments, rats were implanted with the optic fiber (PlexBright optogenetic stimulation system fiber stub implant; 6 mm length) into the vLS, fixed to the skull with two jeweler’s screws and dental cement as described above 21 days after viral transfection and 2 days prior to the experiment. For specific coordinates please see Supplementary Table 4.

### Receptor autoradiography

Brains were cryo-cut into 16-μm coronal sections, slide-mounted, and stored at −20°C. The receptor autoradiography procedure was performed using a linear V1a-R antagonist [125I]-d(CH2)5(Tyr[Me])-AVP (Perkin Elmer, USA) or a linear OXTR antagonist [125I]-d(CH2)5[Tyr(Me)2-Tyr-Nh2]9-OVT (Perkin Elmer, USA) as tracers. Briefly, the slides were thawed and dried at room temperature followed by a short fixation in paraformaldehyde (0.1%). Then slides were washed two times in 50 mM Tris (pH 7.4), exposed to tracer buffer (50 pM tracer, 50 mM Tris, 10 mM MgCl2, 0.01% BSA) for 60 min, and washed four times in Tris buffer 10 mM MgCl2. The slides were then shortly dipped in pure water and dried at room temperature overnight. On the following day, slides were exposed to Biomax MR films for 7-25 days depending on the receptor density and brain region (Kodak, Cedex, France). The films were scanned using an EPSON Perfection V800 Scanner (Epson, Germany). The optical density of V1aR and OXTR binding was measured using ImageJ (V1.37i, National Institute of Health, http://rsb.info.nih.gov/ij/). Receptor density was calculated by sampling the whole region of interest, average gray density was calculated after subtracting the tissue background. Bilateral measurements of 6 to 12 brain sections per region of interest were analyzed for each animal, thus data points represent the mean of those measurements.

### ELISA for plasma corticosterone

Quantification of plasma corticosterone was performed using ELISA. As described before, trunk blood of FIT animals was collected after decapitation. Approximately 1 ml blood was collected in EDTA-coated tubes on ice (Sarstedt, Numbrecht, Germany), centrifuged at 4°C (2000xg, 10 min), aliquoted and stored at −20°C until the assay was performed using a commercially available ELISA kit for corticosterone (IBL International, Hamburg, Germany) following the manufacturer’s protocol.

### Radioimmunoassay for OXT and AVP

OXT and AVP content in extracted blood and CSF, and lyophilized microdialysates was measured using a sensitive and specific radioimmunoassay (RIAgnosis, Germany; sensitivity: 0.3pg/sample cross-reacitivity: <0.7%). All samples were measured within the same assay to avoid inter-assay variability.

### Drugs and viruses

Animals were treated either with endogenous ligands, agonists, or antagonists to modulate the OXT and AVP systems. Usually, drugs were infused 10 min before the FIT in i.c.v. experiments (OXT: 50ng/5μl AVP: 0.1ng or 1ng/5μl, Tocris, Germany). Antagonists were infused 10 min prior to the infusion of the respective agonist with a final volume (agonist+antagonist) of 5μl (OXTR-A, des-Gly-NH2,d(CH2)5[Tyr(Me)2,Thr 4]OVT; V1aR-A, d(CH2)5Tyr(Me)2AVP: 750ng)^37^. For cannulas placed in the LS, agonists (OXT, AVP, TGOT, [Thr4,Gly7]OT: 0.1ng, /0.5μl per side) were infused 5 min before the FIT whereas antagonists (OXTR-A and V1aR-A: 100ng/0.5μl, per side) and muscimol (10ng/0.5μl, per side, Tocris) were infused 10 min before the FIT. In order to modulate the activity of the OXT neurons, the PVN and SON of GH females were infused with 280nl of rAAV1/2 OTprhM3Dq:mCherry (4×10^11^ genomic copies per ml) and rAAV1/2 OXTpr-ChR2:mCherry (4×10^11^ genomic copies per ml) for chemo- and optogenetic experiments, respectively. Viruses were slowly infused by manual pressure infusion at 70 nl/min. After infusion, injected optogenetic animals had their water replaced by salt loaded water for 1 week (2% NaCl) to enhance virus infection and expression. Chemogenetic animals received i.p. infusions either with CNO (HB6149, HelloBio, United Kingdom) or 0.9% saline (1ml/kg).

### Patch-Clamp

#### Slice preparation

Juvenile VGAT-Rats (postnatal days 15-21) were deeply anesthetized with isoflurane and decapitated. Coronal brain slices containing the LS (300 μm) were cut in ice-cold carbogenized (O_2_ [95 %], CO_2_ [5 %]) artificial extracellular fluid (aCSF; [mM]: 125 NaCl, 26 NaHCO_3_, 1.25 NaH_2_PO_4_, 20 Glucose, 2.5 KCl, 1 MgCl_2_, and 2 CaCl_2_) using a vibratome (Vibracut, Leica Biosystems, Germany) followed by incubation in carbogenized aCSF for 30 min at 36°C and then kept at room temperature (~21° C) until experimentation.

#### Electrophysiology

Neurons in the dLS and vLS were visualized by infrared gradient-contrast illumination via an IR filter (Hoya, Tokyo, Japan) and patched with pipettes sized 8–10 MΩ. Somatic whole-cell patch-clamp recordings were performed with an EPC-10 (HEKA, Lambrecht, Germany). Series resistances measured 10-30 MΩ. The intracellular solution contained [mM]: 110 CsCl, 10 HEPES, 4 MgCl_2_, 10 TEA, 10 QX-314, 2.5 Na2ATP, 0.4 NaGTP, 10 NaPhosphocreatine, 2 ascorbate, at pH 7.2. Additionally, biocytin (5 mg/ml) was added to the intracellular solution for post-hoc fluorescent labeling of the patched neurons. The bath aCSF (see above) in the experimental chamber was gassed with carbogen and kept at troom temperature. The average resting potential of lateral septal neurons was −60 mV^42^. Leaky cells with a holding current beyond < −30 pA were rejected. Spontaneous activity (i.e. sIPSCs) was recorded in voltage-clamp mode at resting membrane potential (−60 mV). The frequency and amplitude of sIPSCs were analyzed with Origin 2019 (OriginLab Corporation, Northampton, MA, USA).

#### Histology

After the experiment *in-vitro* slices were post-fixed in 4% paraformaldehyde in PBS (4°C, overnight) and prepared for fluorescent labeling.

### Perfusion

After deep anesthesia with isoflurane followed by CO_2_ rats were transcardially perfused first with 0.1 PBS followed by 4% PFA. Brains were post-fixed in 4%PFA (4°C, overnight).

### Immunohistochemistry

Brains were cut into 40μm slices, which were collected in cryoprotectant solution and stored at −20°C until usage. Typically, a series of 6-8 slices comprehending the whole anteroposterior axis of the region of interest were used for immunostaining. First, slices were washed in 0.1 PBS and then rinsed in Glycine buffer (0.1M in PBS) for 20 min. Afterwards, slices were washed with PBST (0.1 PBS with 0.3% triton-x 100) and blocked for 1 hour in blocking solution. Directly after blocking, slices were incubated in primary antibodies for 1-2h at room temperature and then at 4°C overnight. On the next day, slices were again left in room temperature for 1-2h, washed in PBST and incubated with the secondary antibody. Next, slices were rinsed in 0.1 PBS and mounted on adhesive microscope slides (Superfrost Plus, Thermo Fisher Scientific Inc, USA). Slides were kept in the dark at 4°C until imaging. Especially, for the pERK immunostaining slices were pre-incubate in ice-cold methanol at −20°C for 10 min before the Glycine buffer step. Also for this staining primary antibody incubation lasted 64h. For details on tissue, mounting medium, blocking solution, and antibodies, please see Supplementary Table 5. Imaging from the neural activity, patch-clamp, and molecular identification of the LS neurons was done using an inverted confocal laser scanning microscope (Leica TCS SP8, Leica Microsystems, Wetzlar, Germany). Chemo- and optogenetic imaging was performed using an epi-fluorescence microscope (Thunder Imaging Systems, Leica). Digital images were processed (Merging and Z-projections) using the Leica Application Suite X (Leica) and Fiji^69^. Cell counting was done by an experienced observer blind to the treatments. For patch-clamp, the detailed morphology of the neurites was reconstructed and analyzed from the z-stack with the Fiji plugin Simple Neurite Tracer^70^. From this analysis, the number of branch points, and total branch length of the neurites as well as the area of the soma were extracted and compared between dorsal and ventral neurons of the LS.

### Statistics

Data normality was tested using the Kolmogorov-Smirnov test. If normality was reached, data were analyzed using either Student’s t-test (paired or unpaired), chi-square test, or analyses of variance (one or two way ANOVA) followed by a posthoc comparison corrected with Bonferroni, when appropriate. In case data were not normally distributed, either Mann-Whitney U-test or Dunn’s multiple comparison tests were performed. For detailed statistics information see Supplementary Table 1, 2 and 3.

## Acknowledgments

We would like to thank Rodrigue Maloumby, Anne Pietryga-Krieger, Martina Fuchs, Andrea Havasi, and Thomas Grund for their technical help. VGAT-Venus transgenic rats were generated by Dres Y. Yanagawa, M. Hirabayashi and Y. Kawaguchi in National Institute for Physiological Sciences, Okazaki, Japan, using pCS2-Venus provided by Dr. A. Miyawaki. The OXTR-A and V1aR-A were kindly provided by Maurice Manning (University of Toledo, Toledo, OH). Monoclonal antibodies anti-neurophysin-OXT (p38) and neurophysin-AVP (p41) were kindly provided by Dr. Harold Gainer (NIH, Bethesda). This work was supported by the EU FP7 Project: Neurobiology and Treatment of Adolescent Female with Conduct Disorder: The Central Role of Emotion Processing Fem-NATCD (602407), GRK2174, BO 1958/8-2 to IN. German Research Foundation (DFG) grants LU 2164/1-1 to ML and EG135/3-1, EG135/5-1 to VE. German Research Foundation (DFG) grants GR 3619/7-1, GR 3619/8-1, and GR 3619/13-1, Collaborative Research Center SFB 1158, and Fritz Thyssen Research Foundation (Ref. 10.19.1.015MN) to VG.

## Authors contributions

V.E.M.O. designed, planned, conducted, and analyzed all the experiments, prepared the figures, created the illustrations, and wrote the manuscript. M.L. designed, conducted, and helped with the analyses and interpretation of electrophysiology experiments. H.W., E.D., A.L., A.L.M. conducted and analyzed behavioral experiments. A.B. extracted cerebrospinal fluid. O.B. assisted with surgeries and to conduct optogenetic experiments. V.E. assisted with the design, analyses, and interpretation of electrophysiology experiments. V.G. shared resources, i.e. rAAVs. M.L., V.E., O.B., A.B., and V.G. proof-read and gave input to the manuscript. T.d.J. and I.N. conceived and supervised the project and revised the manuscript.

