## Supplemental Figures 1 to 6 and legends for "Concerted but segregated actions of oxytocin and vasopressin within the ventral and dorsal lateral septum determine female aggression"

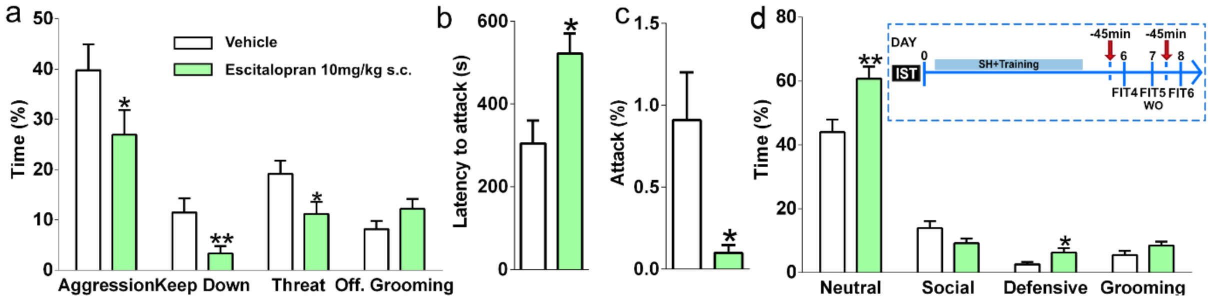


**Supplementary Figure 1. Escitalopram decreases aggression in IST rats. a** Subcutaneous application of escitalopram decreased total aggression (paired two-tailed Student's t-test $t_{(10)}$=2.16, p=0.05), keep down (Mann-Whitney U-test U=26.0, p=0.023), threat ($t_{(10)}$=2.51, p=0.031) and **c** number of attacks (U=24.0, p=0.011) . **b** The latency to attack (U=24.0, p=0.011), and **d** the time spent with neutral ($t_{(10)}$=2.82, p=0.018) as well as defensive behaviors ($t_{(10)}$=2.20, p=0.05) were increased by escitalopram treatment in IST rats. Insert illustrates experimental design (arrow= drug infusions; FIT= female intruder test; IST= isolated and trained; SH= single housing; WO= wash-out). All data are shown as mean+SEM. $^{*}$p<0.05, $^{**}$p<0.01 vs vehicle n=11.

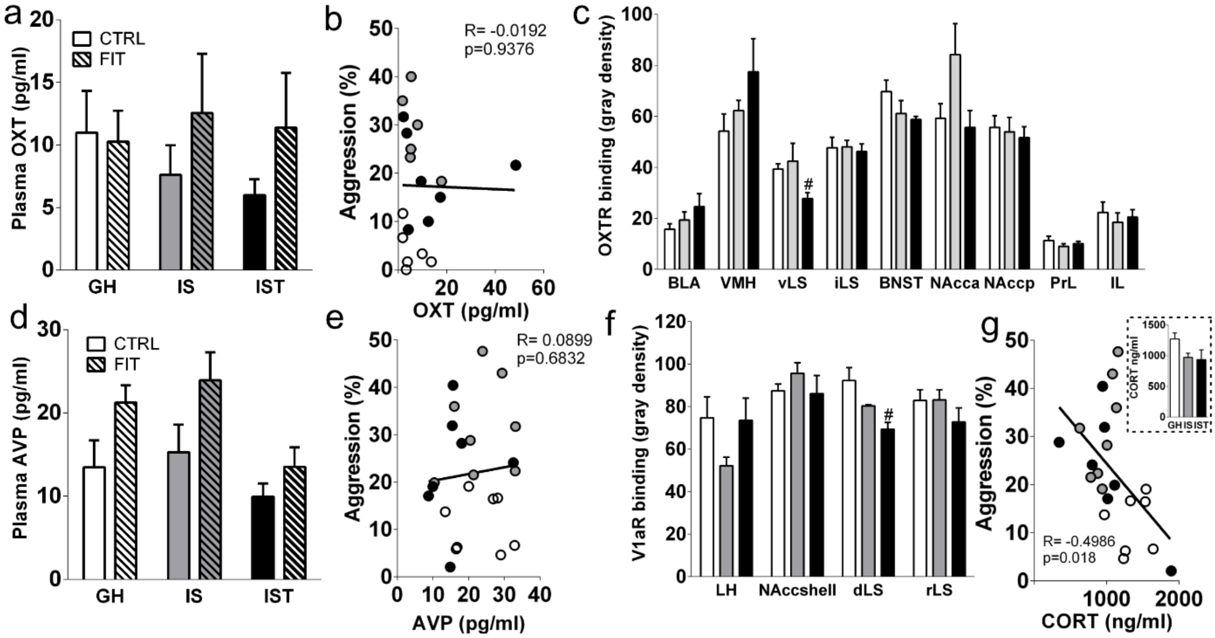


**Supplementary Figure 2. Plasma OXT, AVP, and CORT as well as OXT and V1a receptor binding after exposure to the female intruder test (FIT). a** Plasma OXT remained unchanged in both isolated (IS) and isolated and trained (IST) females in response to FIT exposure, and **b** aggression did not correlate with plasma OXT levels. **c** Among the regions analyzed, OXT receptor (OXTR) binding was only decreased in the ventral lateral septum (vLS) of IST females (Kruskal-Wallis test followed by Dunn's: $H_{(3)}$=7.12, p=0.028). **d** FIT exposure tended to increase plasma AVP levels particularly in group-housed (GH) and IS rats (two-way ANOVA; factor FIT: $F_{(1,55)}$=8.45, p=0.0053; housing: $F_{(2,55)}$=4.39, p=0.017; FIT x housing: $F_{(2,55)}$=0.49, p=0.61). **e** Aggressive behavior did not correlate with plasma AVP concentrations. **f** V1a receptor (V1aR) binding was decreased only in the dorsal LS of IST rats (dLS) ($H_{(3)}$=8.72, p=0.006). **g** FIT exposure did not alter plasma corticosterone (CORT) concentrations (insert). **h** However, aggression negatively correlated with plasma CORT (Pearson's correlation r=-

0.499, p=0.018). All data are presented as mean + SEM. $^{\#}p<0.05$ vs GH. Binding: n=6-9; AVP, OXT: n=7-13; CORT: n=7-8. Abbreviation: basolateral amygdala (BLA), ventromedial hypothalamus (VMH), Infralimbic cortex (IL), intermediate portion of the lateral septum (iLS), bed nucleus of Stria terminalis (BNST), anterior Nucleus accumbens (NAcca), posterior Nucleus accumberns (NAccp), Nucleus accumbens shell (NAcc shell), Lateral hypothalamus (LH), Prelimbic cortex (PrL), rostral portion of the lateral septum (rLS).

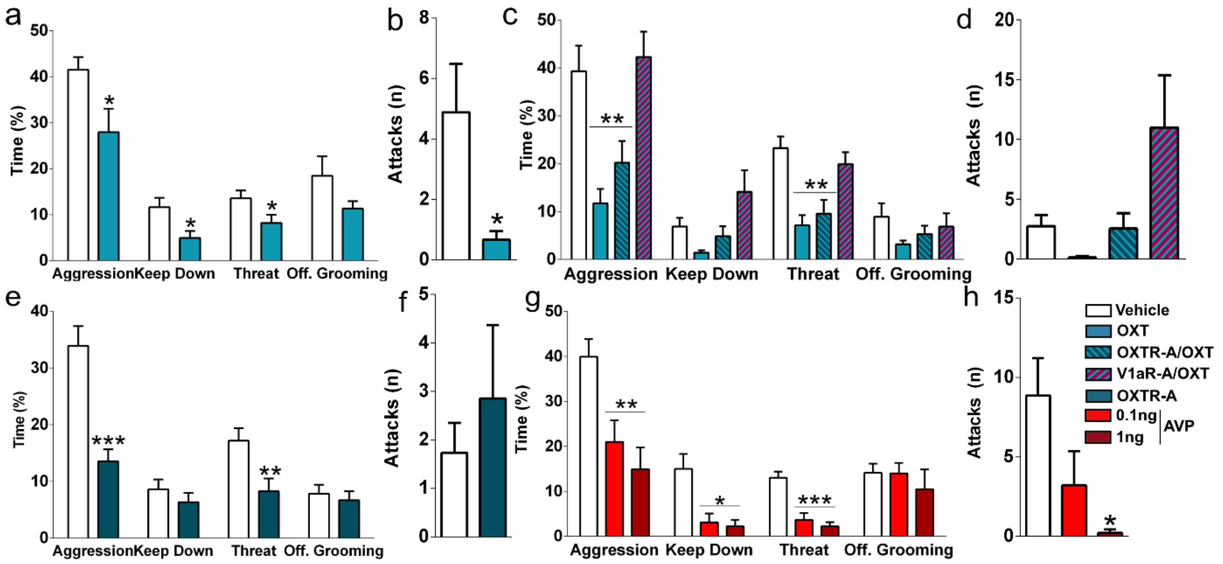


**Supplementary Figure 3. Central activation of OXTRs increases whereas activation of V1aRs decreases aggression in female Wistar rats**. Bar charts depict the effects of intracerebroventricular (i.c.v.) pharmacological manipulations of the oxytocin (OXT) and vasopressin (AVP) systems on female aggression displayed during the FIT by isolated and trained rats. **a** Infusion of synthetic OXT (50ng/5µl) decreased total aggression (paired two-tailed Student's t-test $t_{(8)}=2.33$, p=0.048), keep down ($t_{(8)}=3.17$, p=0.013), threat ($t_{(8)}=1.90$, p=0.045) and **b** number of attacks ($t_{(8)}=2.73$, p=0.026). **c** Infusion of V1aR antagonist (V1aR-A), but not OXTR antagonist (OXTR-A, all at: 750ng/2.5µl) abolished the effects of OXT (50ng/2.5µl) on decreasing total aggressive (one-way ANOVA followed by Bonferroni $F_{(3,28)}=10.08$, p=0.001) and threat behavior ($F_{(3,28)}=9.65$, p=0.0002), **d** without affecting the number of attacks. **e** Blockade of OXTRs by i.c.v. OXTR-A (750ng/5µl) decreased time spent with total aggressive ($t_{(28)}=4.964$, p<0.0001) and threat behavior ($t_{(28)}=2.802$, p<0.0091), **f** but did not affect the number of attacks displayed. **g** Infusion of synthetic AVP (0.1ng and 1ng/5µl) reduced the time spent on total aggression ($F_{(3,54)}=7.483$, p=0.0003), threat (Kruskal-Wallis test followed by Dunn's: $H_{(4)}=25.28$, p<0.0001) and keep down ($H_{(4)}=13.15$, p=0.0043). **h** the higher dose (1ng/5µl) also decreased the number of attacks displayed during the FIT ($H_{(4)}=14.08$, p=0.0028). All data are shown as mean+SEM. $^{*}p<0.05$, $^{**}p<0.01$, $^{***}p<0.001$ or p< 0.0001 vs vehicle. OXT: n=9; AVP: n=9-18; OXTR-A: n=8; Combination OXT/OXTR-A/V1aR-A: n= 7-9.

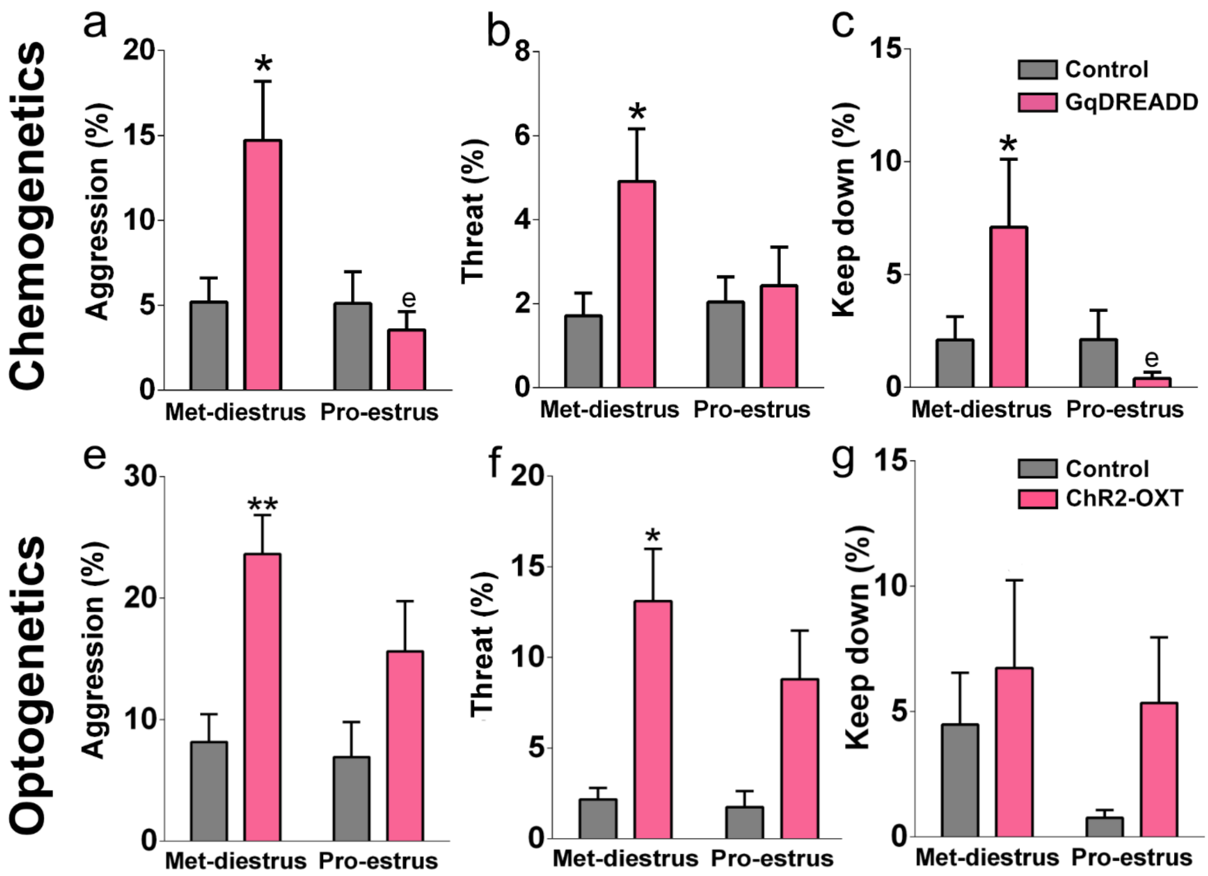


**Supplementary Figure 4. Enhancing central oxytocin (OXT) release by chemogenetic activation of GqDREADD-expressing OXT neurons (a-d) or locally in the ventral lateral septum (vLS) via optogenetic stimulation (e-h) increases aggression in group-housed (GH), non-receptive female Wistar rats. a-c** Administration of clozapine-N-oxide dihydrochloride (CNO, 2mg/kg, i.p.) and subsequent activation of OXT neurons in the paraventricular and supraoptic nuclei of the hypothalamus of females in metestrus and diestrus (met-diestrus) phases of the estrus cycle resulted in increased time spent in **a** total aggressive (factor virus: $F_{(1,19)}=3.34$, p=0.08; estrus cycle: $F_{(1,19)}=6.68$, p=0.018; virus x estrous cycle: $F_{(1,19)}=6.45$, p=0.02), **b** threat (factor virus: $F_{(1,19)}=3.68$, p=0.07; estrus cycle: $F_{(1,19)}=2.96$, p=0.102; virus x estrus cycle: $F_{(1,19)}=5.05$, p=0.037), and **c** keep down (factor virus: $F_{(1,19)}=1.86$, p=0.186; estrus cycle: $F_{(1,19)}=4.94$, p=0.039; virus x estrus cycle: $F_{(1,19)}=4.99$, p=0.038) behaviors. Consequently, GqDREADD rats in proestrus or estrus (pro-estrus) displayed less aggression and keep down compared to met-diestrus rats. **e-g** Blue-light stimulation of OXT terminals in the vLS during exposure to the FIT increased aggressive (factor virus: $F_{(1,15)}=13.06$, p=0.0026; estrus cycle: $F_{(1,15)}= 2.07$, p=0.1708; virus x estrus cycle: $F_{(1,15)}= 1.114$, p=0.308) and threat (factor virus: $F_{(1,15)}=18.52$, p=0.0006; estrus cycle: $F_{(1,15)}= 1.13$, p=0.305; virus x estrus cycle: $F_{(1,15)}= 0.76$, p=0.40) behaviors exclusively in met-diestrus females. All data are shown as mean+SEM. *p<0.05, **p<0.01 vs control; $^{e}$p<0.05 vs met-diestrus. Chemogenetics: n= 7-15; Optogenetics: n= 8-9.

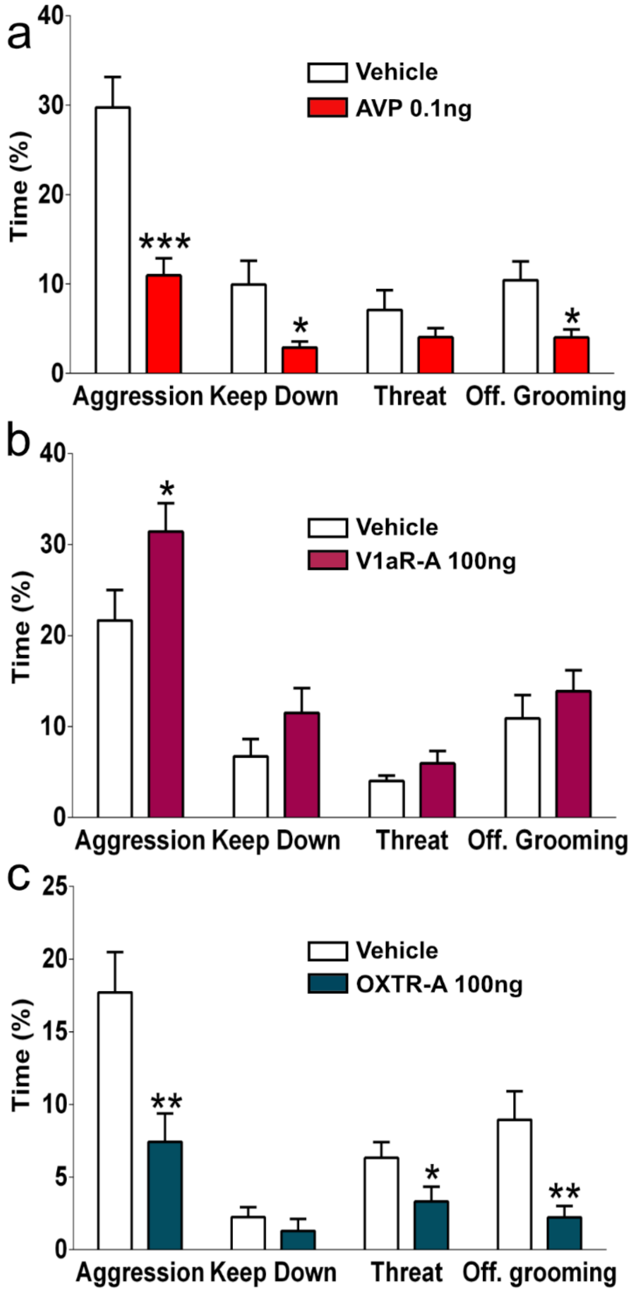


**Supplementary Figure 5. Differential involvement of OXT within the vLS, and of V1a receptors within the dLS on female aggression.** Bilateral AVP infusion into the dLS (0.1ng/0.5µl) decreased the time spent on total aggression ($t_{(22)}$=4.77, p<0.001), keep down ($t_{(21)}$=2.68, p=0.014), threat ($t_{(21)}$=1.3, p=0.21) and offensive grooming ($t_{(21)}$=2.89, p=0.01). **b** Blockade of local V1aRs by infusion of a V1aR antagonist (V1aR-A, 100 ng/0.5µl) into the dLS increased total aggression (two-tailed Student's t-test $t_{(20)}$=2.14, p=0.045). **c** Blockade of local OXTR by infusion of an OXTR antagonist (OXTR-A: 100ng/0.5µl) into the vLS decreased total aggressive ($t_{(26)}$=2.58, p=0.01), threat (Mann-Whitney U-test U=44.0, p=0.024) and offensive grooming (U=44.0, p=0.001) behaviors. All data are shown as mean+SEM. *p<0.05, **p<0.01, ***p<0.001 vs vehicle. AVP: n=12; V1aR-A: n=11-12; OXTR-A: n=13-15.

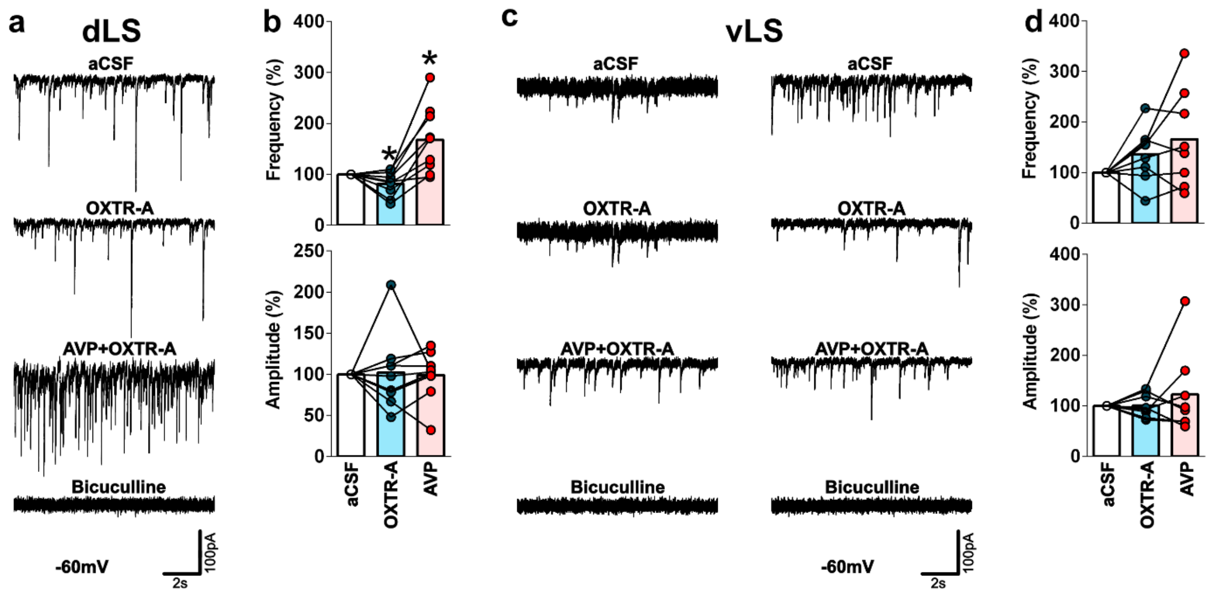


**Supplementary Figure 6. V1aR activation exclusively increases spontaneous inhibitory currents (sIPSCs) within the dLS. a+c** Representative spontaneous current traces during oxytocin receptor antagonist (OXTR-A, 10µM), AVP (1µM) and bicuculline (50µM) bath application in dLS (**a**) and vLS **(c)** cells. **b** OXTR-A decreased sIPSC frequency (Wilcoxon Signed Rank test OXTR-A: $W_{(8)}=-35$, $p=0.04$), whereas AVP increased IPSC frequency ($W_{(8)}=39$, $p=0.02$) in dLS cells. Both treatments did not affect the sIPSC amplitude in those cells. **d** In the vLS cells, neither OXTR-A or AVP had any effect whatsoever on the frequency or amplitude of sIPSCs. $^{*}p<0.05$, vs aCSF. dLS: n=9; vLS: n=8.
